# Phylogeographic evidence that the distribution of cryptic euryhaline species in the *Gambusia punctata* species group in Cuba was shaped by the archipelago geological history

**DOI:** 10.1101/865469

**Authors:** Erik García-Machado, José L. Ponce de Léon, María A. Gutiérrez-Costa, Alice Michel-Salzat, Isabelle Germon, Didier Casane

## Abstract

The main drivers of diversification of freshwater fishes in Cuba are not yet well understood. For example, salt tolerance was thought as the main factor involved in the diversification of *Gambusia punctata* species group in this archipelago. However, evidence from a recent DNA barcoding survey suggested the presence of cryptic species and no correlation between species delimitation and level of salinity. In this study, we analyzed the cryptic diversification of *G. punctata* species group in Cuba, based on a comprehensive sampling of its distribution and including habitats with different salinity levels. We evaluated the patterns of molecular divergence of the samples by sequencing a set of mitochondrial DNA (mtDNA) regions and genotyping nine nuclear microsatellite loci. We also used cytochrome b gene (*cyt*b) partial sequences and these microsatellite loci to analyze population structure inside putative species. Five mtDNA well-differentiated haplogroups were found, four of them also identified by the analysis of the microsatellite polymorphism which corresponds to two already recognized species, *G. punctata*, and *G. rhizophorae*, and three putative new species. The extent of hybrid zones between these groups is also described. In each group, populations inhabiting environments with contrasting salinity levels were identified, indicating a generalized trait not specific to *G. rhizophorae*. The geographic distribution of the groups suggested a strong association with major relict territories of the Cuban Archipelago that was periodically joined or split-up by changes in seawater levels and land uplifts. Salinity tolerance might have facilitated sporadic and long-distance oversea dispersal but did not prevent speciation in the Cuban archipelago.

## 1. Introduction

The complex geological history of the Caribbean archipelagoes has long been considered a major factor promoting population isolation and speciation in different vertebrate groups (Alonso et al., 2012; Glor et al., 2004; Rivas, 1958; Rodríguez et al., 2010; Sly et al., 2011). Among them, the freshwater fishes of Cuba are not the exception (Doadrio et al., 2009; García-Machado et al., 2011; Ponce de León et al., 2014; Rauchenberger, 1989; Rivas, 1958; Rosen and Bailey, 1963). The great diversity of the family Poeciliidae in the Caribbean islands and Continental Middle America (Rosen and Bailey, 1963) makes it a particularly good model to study the history of the Caribbean biotas. Moreover, and in contrast to other freshwater fishes, most species in this family can tolerate a certain degree of salinity, which is thought to be a major barrier to the dispersal of continental species into the islands (Myers, 1938; Rivas, 1958; Rosen and Bailey, 1963). In Cuba, this family is represented by four genera: *Girardinus* (seven species), *Limia* and *Quintana*, with a single species each and *Gambusia* with three species (Fink, 1971; Rauchenberger, 1989; Rivas, 1969). However, recent studies suggested that *Girardinus* (Doadrio et al., 2009) and *Gambusia* (Lara et al., 2010) may be harboring cryptic species. Taxonomy is therefore unclear and refined population genetics and phylogeography studies are needed. Better knowledge about how many cryptic lineages are within these groups and how they are distributed in the Cuban archipelago is crucial to understanding how the family Poeciliidae diversified in the Caribbean and would contribute to the clarification of the history of Middle America biotas.

The estimation of divergence times between fishes in the family Poeciliidae suggested that the ancestors may have colonized the Caribbean islands during the Miocene and Pliocene from different areas of continental America (Reznick et al., 2017). According to Iturralde and MacPhee (1999), during the early Miocene (23-20 Mya), the proto archipelago was characterized by the isolation of landmasses or blocks, due to active tectonic disruption of the Northern and Southern Caribbean plate boundaries. Late in the middle-late Miocene (10-5 Mya), these major land cores (currently Guaniguanico mountains, Isla de la Juventud highs, Guamuaya mountains, and Eastern mountains) corresponding to the current Island of Cuba were progressively connected again by emerging dryland (Iturralde and MacPhee, 1999). Another general uplift occurred during the Pliocene in the region and shaped Cuba as a single territory (Iturralde-Vinent, 2006; van Gestel et al., 1998). During this period, low lands were repeatedly inundated due to sea levels oscillations and earth’s crust movements, producing periods of isolation and ephemeral reconnections (Iturralde-Vinent, 2006). During the last ten thousand years, marine introgressions or desiccation of lakes and rivers due to low precipitations in the Caribbean region have produced recurrent periods of land habitat contractions that should have also impacted local populations (Hodell et al., 2000).

Up to now, very few studies have been dedicated to analyzing the evolutionary history of the freshwater fishes of Cuba using a molecular genetics approach. This is the case for the genera *Girardinus* (Doadrio et al., 2009), *Lucifuga* (García-Machado et al., 2011; Hernández et al., 2016) and *Rivulus* (Ponce de León et al., 2014). The genus *Lucifuga* has a very restricted distribution to hypogean environments of karts patches and the genus *Rivulus* is only present in western Cuba and Isla de la Juventud (formerly Isla de Pinos). *Girardinus* is more widely distributed across Cuba and Isla de la Juventud but it is restricted to freshwater environments (Ponce de León and Rodríguez, 2010; Rivas, 1958; Rodríguez-Machado et al., 2017). Western Cuba seems to represent the diversification center of this genus with only two species present in the Central and Eastern regions (Rivas, 1958; Rodríguez-Machado et al., 2017). The analyses in Doadrio et al. (2009) focused on testing different hypotheses about the origin of this genus rather than interpreting the distribution patterns across the archipelago. The study of other fish species widely distributed across the archipelago may contribute to better understanding the role of its geological history in shaping patterns of species distribution.

In Cuba, there are two species complexes of *Gambusia* fishes: *Gambusia punctata* species complex and *Gambusia puncticulata* species complex. They both are widely distributed across Cuba and Isla de la Juventud (Ponce de León and Rodríguez, 2010), which make them excellent models to assess their diversification in the Cuban archipelago using genetic markers. In the present study, we analyzed the genetic diversity within the *Gambusia punctata* species complex.

The *Gambusia punctata* species group contains five species (Rauchenberger, 1989): *Gambusia punctata* (Poey 1854) endemic to Cuba, *G. rhizophorae* (Rivas 1969) found in northwestern Cuba and south Florida including the keys, *G. xanthosoma* (Greenfield 1983) from Grand Cayman Islands, *G. beebei* (Myers 1935) and *G. pseudopunctata* (Rivas 1969) both endemic to Haiti. According to Rivas (1969), *G. punctata* and *G. rhizophorae* show differences in body shape and in the frequency of a number of gonopodial characters. However, most of the meristic character measured actually overlap between the two species (e.g. number of dorsal fin rays, number of lateral body scales, number of segments distal to gonopodium elbow, see Rivas (1969) for details), making difficult the identification of individuals following these criteria. Rivas (1969) reported that these two species have separated distributions according to differences in salinity tolerance; *G. punctata* is widely distributed in most Cuban island and Isla de la Juventud but restricted to freshwater habitats, while *G. rhizophorae* inhabits brackish and salt water, and is distributed along the north-western coast of Cuba and coastal wetlands of South Florida (Southeast US) (see Rivas 1969, Fig. 5). Other early works have mentioned other Cuban “forms” within the *G. punctata* species group, from east-central-localities (i.e. *G. punctata punctulata* from Remedios, *G. filamentosa* from Viana River in “Sagua la Grande” and *G. finlayi* from Camagüey Province), but descriptions in all these cases were vague and subsequent analysis did not show morphological divergences supporting them (Rivas, 1969). Recently, Rodríguez (2015a) reported new distribution records for *G. rhizophorae* in brackish or saltwater habitats along the north-eastern and south-eastern coasts of the Cuban archipelago basing the identification of the specimens on the criteria of Rivas (1969). However, it is not clear which of the variables determined by Rivas (1969) were used to assure the correct identification of the species. Also, neither the variation in the analyzed characters nor the sample sizes used for analysis are shown.

Based on a barcoding DNA analysis (Lara et al., 2010), a third Cuban cryptic lineage (*Gambusia* sp.) was found within *G. punctata* species group. It contains individuals collected at two localities, a brackish sinkhole (La Jenifer) at Key Coco, north-central region, and Yara River in Guantánamo province, in the eastern region. The estimates of genetic divergence (using Kimura’s two parameters model, thereafter K2P model) between the three Cuban lineages ranged between 4.9 ± 0.8% between *G. punctata* and *Gambusia* sp., and 5.2 ± 0.8% between *G. punctata* and *G. rhizophorae*, in all cases exceeding the 2% cut-off used to identify freshwater fish species (April et al., 2011; Pereira et al., 2013) and the 3% cut-off applied to separate other vertebrate species (e.g. Vieites et al., 2009). However, mitochondrial DNA divergence as a yardstick for species delimitation has been strongly criticized by different authors (Moritz and Cicero, 2004) and poor prediction of nuclear gene divergences suggests that mtDNA divergence is not sufficient to delimitate species (Hutson and Turelli 2003). Additionally, such genetic distance thresholds must be supported by an adequate sampling of intraspecific diversity and comparison of sister species (Sites & Marshal 2003; Johnson and Cicero 2004). That said, the results of the barcoding analysis opened new questions about taxonomy and speciation processes in this group. Indeed, the geographic and ecological distribution of *Gambusia* sp. raised doubts about variations in salinity tolerance as the main driver of the divergence between *G. punctata* and *G. rhizophorae* proposed by Rivas (1969), as well as the occurrence of *G. punctata* in east-central territories of Cuba. Moreover, salinity tolerance makes *Gambusia* fishes particularly interesting to contrast patterns observed in other terrestrial species highly dependent on land connections to disperse. If dispersal across marine barriers was common during the evolutionary history of *Gambusia* species in Cuba, one can expect a relative genetic homogeneity among populations distributed over long distances. However, if historical disconnections among land cores shaped evolutionary divergence of different populations and speciation, one can expect a correspondence between the geographic distribution of genetic groups and subaerial land cores. The aim of the present study was to analyze the genetic diversification of *G. punctata* species complex in Cuba, on the basis of a comprehensive sampling of its distribution including saltwater, brackish and freshwater habitats, in order to test the hypothesis that the geological history of the Cuban archipelago is the main factor which drove the divergence and the distribution of the main groups. We sequenced a set of mtDNA regions and genotyped nine nuclear microsatellite loci in order to describe and analyze the pattern of genetic differentiation according to the geographic distribution of the samples. Five main mtDNA haplogroups were found, two more than the three previously identified (Lara et al., 2010). These haplogroups are geographically isolated but not associated with different salinity levels. Four of these groups were also supported by the analysis of the nuclear DNA polymorphism. Their geographic distribution suggested a strong association with major relict territories of the Cuban archipelago that were episodically joined or split-up by changes in seawater level and land uplifts. We also used *cyt*b partial sequences and these microsatellite loci to decipher genetic structures within these groups. This study contributes to a much better understanding of several aspects of the phylogeography of the *Gambusia punctata* species group in Cuba and the Caribbean.

## 2. Materials and methods

### 2.1 Samples and localities

In order to have a comprehensive representation of the genetic diversity of the *G. punctata* species group, including fresh, brackish and salt-water environments, we sampled most of the distribution area in Cuba (Fig. 1; Supplementary material 1a). A total of three hundred four (n = 304) fishes were collected at 66 localities using hand nets or seine nets depending on the river depth and topology. Fish capture and sample collections were done under the permits: CH-40-DB (026) 08 and CH-8116247-5, issued by the Cuban Centre for Environmental Inspection and Control (CICA). Prior to fixation in 95% ethanol, the fish were euthanized with tricaine meta-sulphonate (MS-222). All specimens are preserved in the collection of the Centro de Investigaciones Marinas, University of Havana.

**Figure 1.**
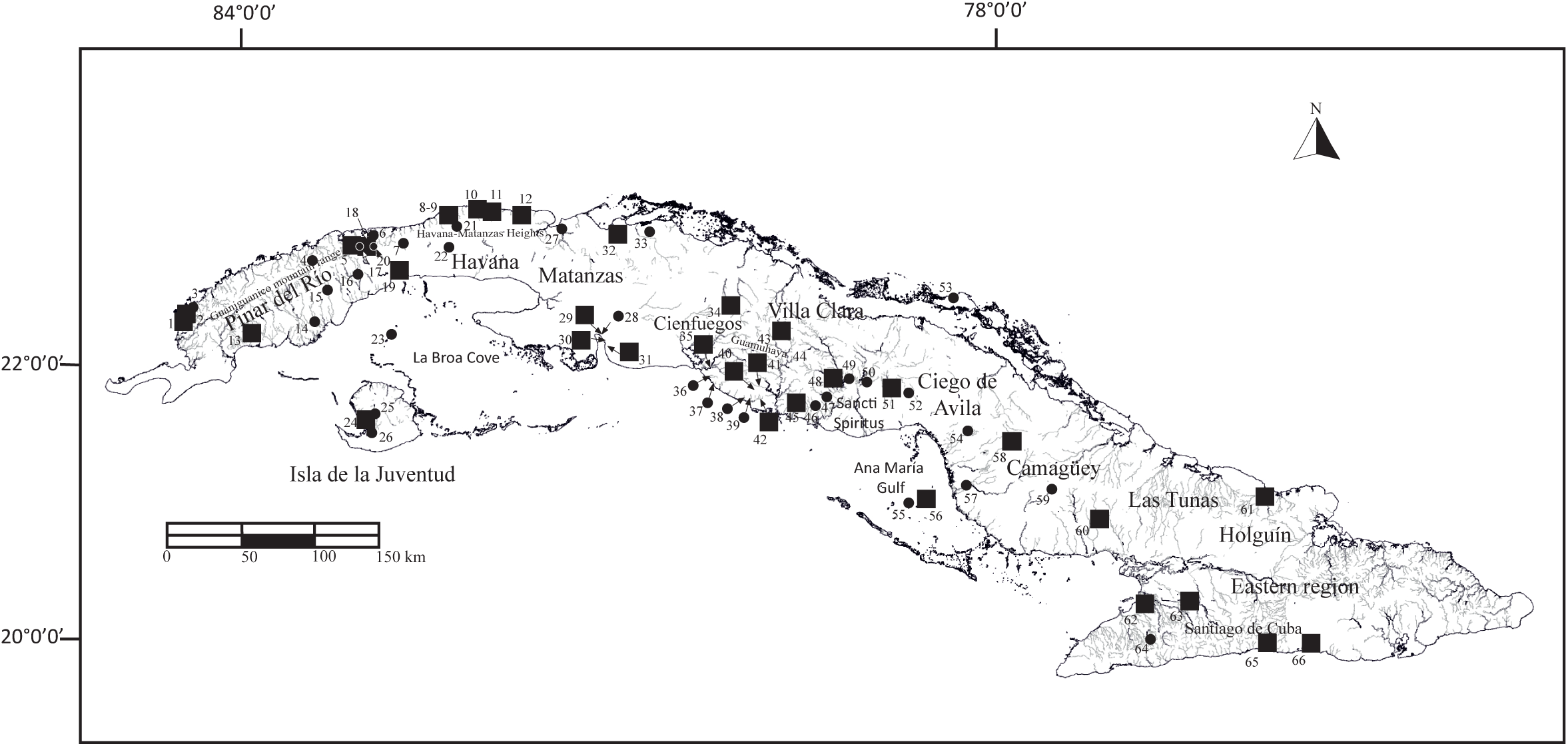
Sampling sites for *Gambusia punctata* species group along the Cuban archipelago. Black circles indicate sites where samples were analyzed with *cyt*b sequences alone, and black boxes indicate sites where samples were analyzed with *cyt*b sequences and microsatellites. Locality numbers are in reference to Supplementary material 1a.

### 2.2 DNA extraction, mtDNA amplification, and analysis

We extracted DNA from small portions of muscle tissue (~3mm) using the NucleoSpin® Tissue Kit. For a total of 141 individuals, the 5’ domain (752 bp) of *cyt*b was amplified using polymerase chain reaction (PCR) with primers GluGamb (5’ACT CAA CTA TAA GAA CYC TAA TGG C) modified from Meyer et al. (1990) and CB3 (5’ TGC GAA GAG GAA GTA CCA TTC) (Palumbi, 1996). A subset of specimens (n = 22), standing for the different *cyt*b lineages, were additionally amplified for six other mtDNA regions: 12SRNA - 16SRNA, 16SRNA – tRNALeu – ND1, COI, COII-tRNALys, COIII and control region that represented, including the *cyt*b, a total of 4,763 nucleotide sites (Supplementary material 1 a and b) to increase resolution of the mtDNA phylogenetic analysis. The polymerase chain reaction (PCR) was performed with 5-100 ng of total DNA in a 50 µL final reaction volume. The reaction contained 1 unit of GoTaq DNA polymerase (Promega), 1X enzyme Flexi Buffer, 0.25 µM of each primer, 0.2 mM dNTPs, and 2 mM MgCl2. The PCR products were purified using the Illustra ExoStar 1-Step kit (General Electric Company), and a volume of 0.8 µL (5-30 ng) of the purified product was used for both sides sequencing with the Big Dye terminator sequencing kit (Applied Biosystems). The fragments were resolved in an ABI 3100 automated sequencer (Applied Biosystems). The sequences were deposited in the EMBL database under the accession numbers provided in Supplementary material 1a. The sequences of *G. puncticulata* C (*sensu* Lara et al., 2010) amplified and sequenced with the same set of primers were used as outgroup. In the case of the *cyt*b, available sequences of *G. punctata* (U18220), *G. rhizophorae* (U18223) and *G. hispaniolae* (U18209) (Lydeard et al., 1995), and *G. rhizophorae* (KM658368) (Heinen-Kay et al., 2014) were also included in phylogenetic analyses.

Previous to alignment, the raw sequences were inspected by eye against the chromatogram using BioEdit Sequence Alignment Editor v7.0.8.0 (Hall, 1999). The alignments were performed with Clustal W (Thompson et al., 1994) in MEGA v7.0.26 (Kumar et al., 2016). The seven gene regions amplified from selected individuals were concatenated and considered as a single unit for the analysis. The program MEGA v7.0.26 was also used to infer the model best fitting the nucleotide substitution parameters of the data set, using a Neighbor-joining tree and a strong branch swap filter option. The model was selected using the Bayesian information criteria (BIC). We used maximum likelihood (ML) and Bayesian methods to infer phylogenetic relationships. The phylogenetic trees were obtained for the *cyt*b and the concatenated alignment sets. The ML phylogeny was obtained using MEGA v7.0.26 (Kumar et al., 2016). The parameters of the selected model, nucleotide substitution matrix, gamma-distributed rate variation across sites, and proportion of invariant positions were used for tree inference. Stationary base frequencies and substitution rates were optimized during tree inference. The ML tree was found by heuristic search from an initial tree obtained by Neighbor-Joining and BioNJ algorithms, optimized with the Nearest-Neighbour-Interchange (NNI) algorithm and a very strong branch swap filter for an exhaustive search. The robustness of the nodes of the ML tree was assessed using the bootstrap method with 1,000 replicates.

For Bayesian tree reconstruction with MrBayes 3.2 (Ronquist et al., 2012), model parameters (*i.e.* substitution rate matrix and stationary nucleotide frequencies) were those inferred from ML analyses. The Bayesian inference was initially based on two independent runs using four Metropolis-coupled Monte Carlo Markov chain for 2 × 10^6^ generations, with sampling every 200 generations. Convergence was confirmed examining various diagnostic outputs, particularly the Potential Scale Reduction Factor (PSRF) and the Effective Sample Size (ESS). The ESS values were obtained using TRACER v 1.6 (http://beast.bio.ed.ac.uk/). The MCMC chains were then run for a further 4 × 10^6^ generations. The PSRF value approached one (PSRF = 1.002) and the ESS values were greater than 200, ranging from 1105.84-15025.75. The first 25 percent of the sampled trees were discarded as burn-in. The sampled trees from both runs were used to construct a consensus tree and to estimate the posterior probabilities of the nodes.

The relationships between haplotypes within each group were visualized as networks inferred using the Median-Joining network algorithm (Bandelt et al., 1999) and the Maximum Parsimony (MP) option to remove superfluous median vectors and links not present in the shortest trees (Polzin and Daneshmand, 2003) which are implemented in Network 5.0.0.1 (Fluxus-engineering.com). Defaults settings were used in all cases.

To infer putative species boundaries, we used the Poisson tree processes (PTP) model (Zhang et al., 2013). Based on the phylogenetic species concept, this method models the speciation rates directly from the number of substitutions under the assumption that the number of substitutions is higher between than within species. We used the version implemented in the web site (https://species.h-its.org/ptp/). This performs a Bayesian implementation (bPTP) of the model and provided Bayesian support for the putative species boundaries. The combined and the *cyt*b trees obtained with the program RAxML (Stamatakis, 2006) were used as inputs. The RAxML phylogeny was conducted on an input alignment pruned of identical haplotypes and using the GTRCAT substitution model and the Hill-climbing algorithm (Stamatakis et al., 2007) for a heuristic tree searching from a predefined initial tree and constraining the outgroup to obtain a rooted tree. The number of MCMC generations for bPTP was set to 500,000 while the other parameters were used as predefined (thinning = 100; burnin = 0.1).

Sequence divergence within and between groups was estimated using *p* distances. The variance of the estimates was obtained by 1,000 bootstrap repetitions. All estimates were obtained using MEGA v7.0.26 (Kumar et al., 2016).

### 2.3 Microsatellite loci amplification and analysis

We used nuclear loci to contrast the hypothesis obtained by analyzing the mtDNA sequences. Nine microsatellites loci were amplified from a total of 228 individuals from 33 localities encompassing the geographic distribution of the species group. The loci analyzed were: Gaaf10, Gaaf13, Gaaf15, Gaaf16, Gaaf22 (Purcell et al., 2011), Gafµ5 (Spencer et al., 1999), Mf6, Mf13 (Zane et al., 1999), GG2B (Cureton et al., 2010). The PCR reactions were carried out in 10 µL of final volume with the following reaction conditions: 5 - 20 ng of total DNA, 1X Platinum^TM^ Multiplex PCR Master Mix (ThermoFisher Scientific), and 0.2 µM of each marked primer. The program used was 2 min at 95 °C, followed by 30 cycles of 30 s at 95 °C, 90 s at 60 °C and 60 s at 72 °C, and a final extension for 30 min at 60 °C. Genotypes were scored using an ABI 3130 XL Genetic Analyzer with GS500(−250) LIZ size standard and the software Genemapper 3.0 (Applied Biosystems).

The presence of microsatellite null alleles was tested using Micro-Checker 2.2.3 (Van Oosterhout et al., 2004) and Hardy-Weinberg equilibrium was assessed by estimating the exact probability of the *F*_IS_ statistic Weir and Cockerham (1984) using the Markov chain method in GENEPOP 4.7 (Rousset, 2008). The pairwise loci linkage disequilibrium was evaluated using the log-likelihood ratio *G*-statistic in FSTAT 2.9.3 (Goudet, 2001).

In order to determine the number of genetic clusters defined by the microsatellite loci, we used the software STRUCTURE 2.3.3 (Pritchard et al., 2000). The method probabilistically assigns individuals to one or more populations based on their genotypes providing a level of admixture in each case. The “admixture model” and allele frequencies correlated among populations were used as priors to cluster individuals according to their shared population ancestry. Twenty runs were performed for each number (*K*) of clusters tested. The run-length was set to 10^6^ MCMC (Markov chain Monte Carlo) repetitions after a burn-in period of 5 × 10^5^. Evanno’s method (Evanno et al., 2005) was used to estimate the best value of *K* as implemented in STRUCTURE HARVESTER v.0.6 (Earl and vonHoldt, 2012). CLUMPP version 1.1.2 (Jakobsson and Rosenberg, 2007) was then used to align individual posterior assignment probabilities from independent replicates.

The D statistic of Jost (2008) corrected for sampling bias was used to estimate divergence among groups. This statistic is based on the effective number of alleles and is expected to better estimate divergence among populations when the number of populations is small (Jost, 2009; Jost et al., 2018; Meirmans and Hedrick, 2011). The statistical significance and confidence intervals were obtained by jackknifing using GenoDive v 2.0b27 (Meirmans and Van Tienderen, 2004). The fixation index (*F*_ST_) statistic of Weir and Cockerham (1984) was also estimated to compare with estimations recorded from other species pairs. The distribution of pairwise *F*_ST_ estimates under the hypothesis of no difference between two clusters was obtained by permuting 10,000 times genotypes between clusters using Arlequin 3.5.1.2 (Excoffier and Lischer, 2010).

### 2.4 Population subdivision inside groups

We used analysis of molecular variance (AMOVA) (Excoffier et al., 1992) on *cyt*b sequences to test for possible population partitions inside groups based on results of haplotype distributions and the different geographic regions (e.g. plains, mountains, island, and keys) inside area distributions. Significance for the different hierarchical level of partition variance was obtained with 16,000 permutations. STRUCTURE analysis (Pritchard et al., 2000) was conducted independently for each group following the same procedures described above. The number of *K* partitions tested varied according to the geographic distribution of the group.

### 2.5 Genetic diversity and demography

Within each group, haplotype diversity (*Hd*) and nucleotide diversity (π) of the *cyt*b sequences were estimated using DnaSP v. 5.10 (Librado and Rozas, 2009). For the microsatellite loci, the percentage of polymorphic loci, the number of alleles and private alleles, and the observed (*Ho*) and expected (*He*) heterozygosities were calculated using FSTAT 2.9.3 (Goudet, 2001). To test for demographic changes (*i.e*. population growth), Tajima’s *D* (Tajima, 1989) and Fu’s *Fs* (Fu, 1997) neutrality statistics, which are sensitive to demographic changes, were performed. Tajima’s *D* test is based on the expectation that under mutation-drift equilibrium, *θ* (Watterson, 1975) and π should give the same estimation of 4*Nu*, and thus significant differences may indicate a departure from neutrality. Fu’s *Fs* is based on the contrast between the number of haplotypes and the number of samples drawn from a constant-sized population. Under the effect of selection and hitchhiking it is expected that both statistics tend to departure from expected proportions. However, under population growth, *Fs* is more powerful and should show significant negative values. The statistics and their statistical significance, estimated by 10,000 coalescence simulations, were obtained using DnaSP v. 5.10 (Librado and Rozas, 2009).

## 3. Results

### 3.1 Phylogenetic relationships using mtDNA sequences

A total of 141 *cyt*b sequences were obtained from individual fish of samples which are representative of the distribution area of the *G. punctata* species group (Fig. 1, Supplementary material 1a). The results of the phylogenetic analysis using maximum likelihood and Bayesian methods are presented in Fig. 2A (detailed trees in Supplementary material 2). The tree reconstruction was based on a 752 bp sequence, using the TN93 (Tamura and Nei, 1993) nucleotide substitution model with a gamma distribution of rates across sites (α = 0.14) and invariant sites (I = 0.58). Further details on parameters are provided in Supplementary material 2a and 2c. Tree topology was highly congruent with both methods. A tree constructed with seven mtDNA partial gene sequences (*cyt*b, 12SRNA - 16SRNA, 16SRNA – tRNAIle –ND1, COI, COII-tRNALys, COIII, and CR) totaling 4,763 bp, had similar topology for the three internal nodes but with stronger support (Supplementary material 2b). Four major mtDNA haplogroups were identified. The first one, identified as *G. rhizophorae*, is distributed along the northwestern region from Pinar del Río to eastern Havana as recognized from the original description (Fig. 2A, C). *G. rhizophorae* populations are geographically restricted to Guaniguanico mountain range and Havana - Matanzas Heights with populations in fresh and brackish waters (Fig. 2C, Supplementary material 1). Interestingly, the sequences of *G. rhizophorae* from Florida (GenBank ascension numbers U18223 and KM658368) are nested within this clade (Supplementary material 2a). Individuals from Mil Cumbres represent a divergent sub-haplogroup within this clade (site 4 in Fig. 2C). The second haplogroup, including GenBank sequence U18220 (Lydeard et al., 1995), was not recovered by either method with the *cyt*b only but received strong support with the concatenated data set (Supplementary material 2). It is distributed from southwestern Pinar del Río to Matanzas provinces and Isla de la Juventud and was identified as *G. punctata*. Its populations were found in rivers from the western plains and mountain rivers/streams (i.e. localities 17; 18; 20; 24; 25) of the region. Some populations were found in brackish and saltwater (19; 23; 26). Both *G. rhizophorae* and *G. punctata* haplotypes were sympatric in Baracoa River (Fig. 2A, C). Four *cyt*b sequences were identified as *G. punctata* and 11 as *G. rhizophorae*, and both species were syntopic at localities 8 and 9 (Fig. 1, Supplementary material 1a). The third haplogroup, named here *Gambusia* sp. according to Lara et al. (2010) is distributed from eastern Villa Clara to Santiago de Cuba (Centro-Eastern region) provinces. Populations in saltwater were found in keys at the north (53) and the south (55; 56). The fourth haplogroup is distributed from eastern Cienfuegos to Villa Clara, in Guamuhaya mountain range and surrounding areas and defined as *Gambusia* sp. D (i.e. to follow Lara *et al*., 2010 designations). Noteworthy, one population was identified in Canimar River (27), North Matanzas province, inside *G. punctata* distribution area (Fig 2A, C). Although sampled in a freshwater source in the riverbank, it might be systematically exposed to salinity level variations due to its proximity to the river mouth.

**Figure 2.**
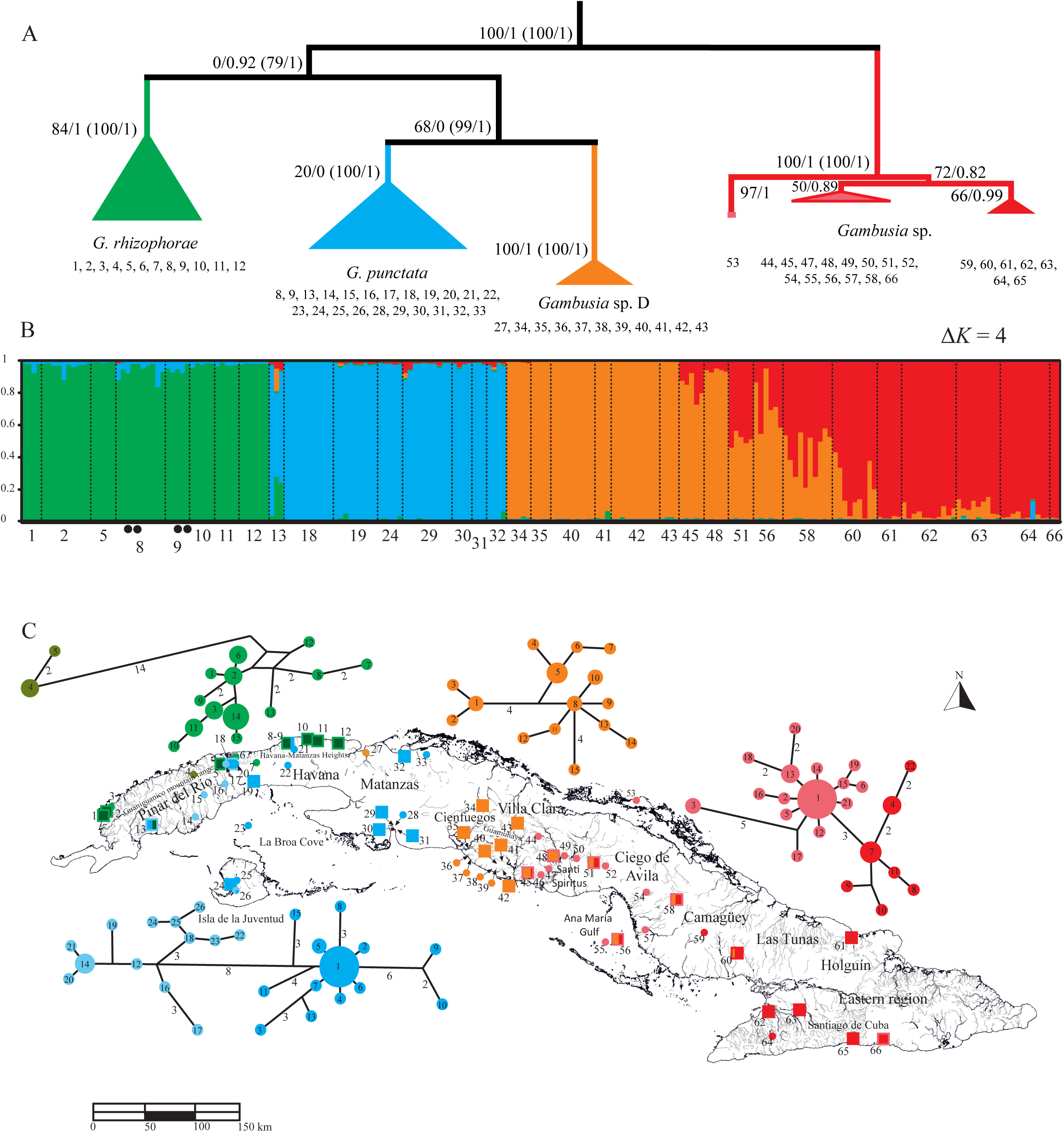
Maximum likelihood phylogenetic tree based on 752 bp *cyt*b sequences (A); STRUCTURE analysis based on 9 microsatellite loci (B) and *cyt*b haplotype networks for each group and map showing group distributions (C). Colors distinguish the four groups identified by the mtDNA and microsatellite datasets and phylogenetic and STRUCTURE analysis: Green (*Gambusia rhizophorae*); blue (*G. punctata*); orange (*Gambusia* sp. D); red (*Gambusia* sp.). Numbers below figures (A) and (B) represent sampling locality numbers as in map (C) and Supplementary material 1a. (A) The *cyt*b tree is showed compressed for each group, and external taxa used for rooting the tree were pruned. ML and Bayesian trees were obtained using the Tamura-Nei (1992) model with a gamma distribution (α = 0.14) and invariants positions (I = 0.58). Values beside branches indicate bootstrap and Bayesian posterior probabilities of nodes respectively. Between parenthesis are the corresponding values of bootstrap and Bayesian posterior probabilities obtained with the concatenated sequence set. (B) STRUCTURE graphic of individual membership probabilities obtained after Evanno’s test (Δ*K* = 4). Black dots on the bottom of the graphic indicate individuals with *G. punctata* haplotypes. (C) Haplotype networks are portrayed above or below the map according to the geographic distribution. Haplotype numbers are inside circles (see Supplementary material 1a for distribution). Boxes and circles are as in Fig. 1. Outline and inner colors correspond to mtDNA and microsatellite results respectively.

The tree constructed with concatenated mtDNA sequences strongly supports the sister relationship between *G. punctata* and *Gambusia* sp. D. while the node relating *G. rhizophorae* as sister to this clade was only well supported by Bayesian inference (Bayesian posterior probability = 1; ML bootstrap = 79%) (Supplementary material 2b). *Gambusia* sp. was the first group to split off. The putative species boundaries using the bPTP model analysis on the *cyt*b and the combined sequence set agree in suggesting most main groups (Supplementary material 3). As expected, the combined sequence set outperformed the *cyt*b results on this analysis (see Zhang et al. 2013). However, we only observed strong support for *Gambusia* sp. with the concatenated data set (Bayesian support = 0.976). Interesting, the bPTP analysis with the combined sequence set provides some support for a split of *G. rhizophorae* into two separate groups, the first including most of the populations and the second, including Mil Cumbres population only (Supplementary material 3).

With *cyt*b, mean uncorrected *p* distances within haplogroups were relatively low ranging between 0.4% ± 0.12 in *Gambusia* sp. and 1.6% ± 0.19 in *G. rhizophorae* (Table 1). In this latter case, when split into the two groups defined above, the estimates were much lower: 0.1% ± 0.1 in Mil Cumbres and 0.4% ± 0.1 in *G. rhizophorae* without Mil Cumbres). In contrast, mean sequence divergences between haplogroups were high in all pairwise comparisons ranging from 3.4% ± 0.6 between *G. punctata* and *Gambusia* sp. D to 5.5% ± 0.7 between *Gambusia* sp. D and *Gambusia* sp. The mean distance between Mil Cumbres and other *G. rhizophorae* populations was also high (2.7% ± 0.5).

**Table 1.**
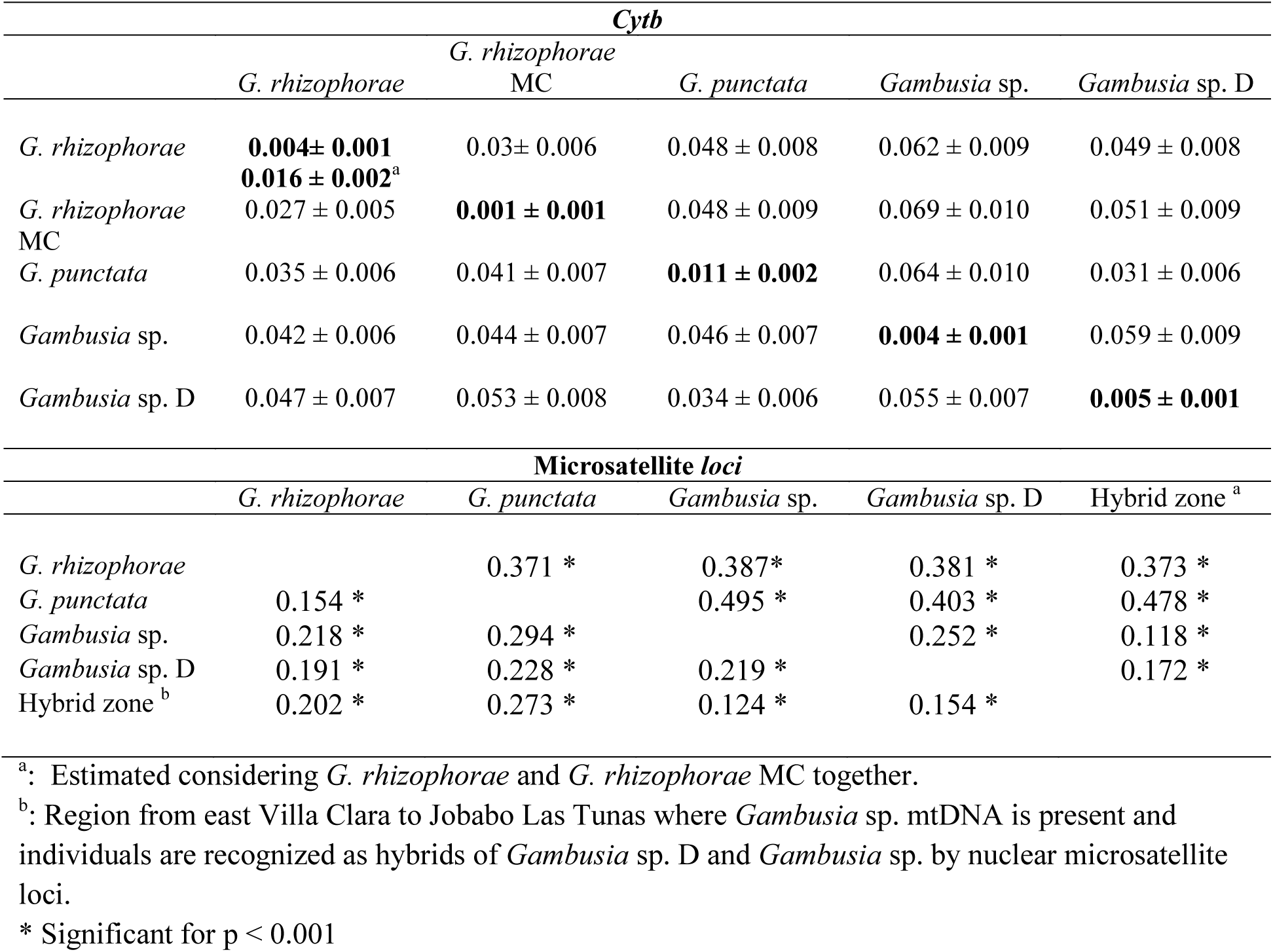
Divergence estimates (*p* distances) between the different lineages using *cyt*b above the diagonal and COI sequences below the diagonal. Within *cyt*b haplogroup divergence estimates are on the diagonal. Microsatellite *loci* Weir and Cockerham (1984) *F*_ST_ estimates are below the diagonal and Jost (2008) D estimates are above the diagonal.

### 3.2 Cluster identification using microsatellite loci

Of the nine microsatellite loci, null alleles were detected at four of the 33 analyzed localities. Null alleles were present in *Gaaf* 10, *Gaaf* 13, *Gaaf* 15, and *Gaaf* 16. No departure from the expected random allele combinations between the different *loci* was found at the whole data set level (Bonferroni correction for multiple tests, α = 0.00139), nor at the population level (Bonferroni correction, α = 0.000046). At all loci and at most localities, genotypes were in Hardy-Weinberg proportions. After Bonferroni correction, only 3 loci showed heterozygote deficiency (Gaaf16 at Camarones River, Gaaf22 at Cojimar River, and Gaaf13 at Bayamo River). Given that no locus showed recurrent bias from expected proportions, all loci were used in subsequent analysis. For some loci, a single allele was found at several localities (Supplementary material 1 and 4).

The Bayesian clustering of 228 individuals in 2 to 7 partitions using STRUCTURE (Pritchard et al., 2000) supported recognition of four genetic clusters (Fig. 2B) using Evanno’s test (Supplementary material 5a). Using this method, the species *Gambusia rhizophorae* and *G. punctata* were well delimited, as they were in the phylogeny obtained with mtDNA sequences. However, the geographic distribution of microsatellite genotypes in *Gambusia* sp. D and *Gambusia* sp. presented a wide region of genetic intergradations that extend from east Villa Clara to Las Tunas (hereafter hybrid zone) where membership probabilities of individuals gradually change from western (*Gambusia* sp. D) to eastern (*Gambusia* sp.). The STRUCTURE analysis showed complete differentiation between western and eastern populations (Fig. 2B, C). In contrast with the results from mDNA, the pattern found through the analysis of microsatellite loci indicates directional introgression with a complete occurrence of *Gambusia* sp. mtDNA across the hybrid zone (see localities 45, 48, 51, 58, 60) (Fig. 2C). In order to estimate genetic differentiation between the four putative species, we excluded individuals from the hybrid zone that is with the highest membership probability below 90%, sampled from Calabazas River (44) to Jobabo River (60). The genetic differentiation (*F*_*ST*_) estimated between pairs of clusters was statistically significant (p < 0.001) in all cases, ranging from 0.154 [*G. rhizophorae* / *G. punctata*] to 0.294 [*G. punctata* / *Gambusia* sp.] (Table 1). Interestingly, significant *F*_*ST*_ were found between the hybrid zone and its parental groups: 0.124 and 0.154 with *Gambusia* sp. and *Gambusia* sp. D, respectively. These *F*st values represent about half of the value (*F*st = 0.219) estimated between the parental species.

### 3.3 *Analysis of sympatry of G. rhizophorae* and *G. punctata in Baracoa River*

As mentioned above, *G. rhizophorae* and *G. punctata* haplotypes are sympatric in Baracoa River. However, microsatellite loci analysis indicated that four individuals having *G. punctata* haplotypes were clustered confidently as *G. rhizophorae* (Fig. 2B). To perform a more detailed analysis on this finding, the number of samples was increased from 15, in the first general analysis, to 31 including all individuals from Baracoa River B1 and B2 localities (also localities 8 and 9 in Supplementary material 1a). The full set of samples from other localities belonging to both species was also included. Three more individuals with *G. punctata cyt*b haplotypes were identified totaling seven individuals with *G. punctata* mtDNA within the *G. rhizophorae* cluster (Fig. 3A). Interestingly, the STRUCTURE analysis of microsatellite loci also revealed some degree of hybridization at Papaya River. This location is close to the western limit within the distribution area of *G. punctata*. Two of the three individuals analyzed from this locality showed shared membership probabilities to both *G. rhizophorae* and *G. punctata* (Fig. 3A).

**Figure 3.**
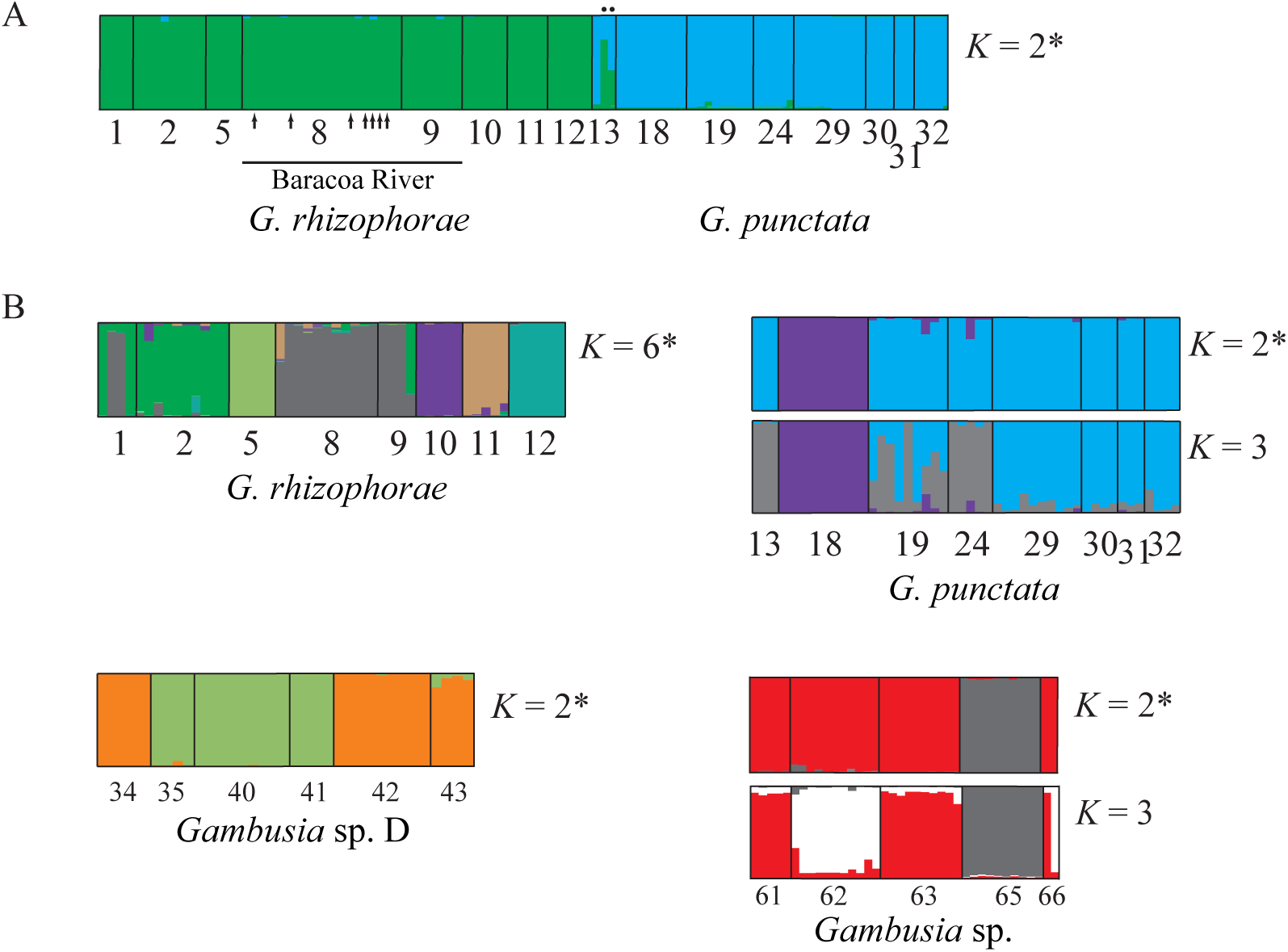
STRUCTURE graphics showing the results of the analysis with different data sets and *K* values. The *K* determined after Evanno’s test are indicated by an asterisk. (A) Analysis of *Gambusia rhizophorae* and *G. punctata* with the larger sample size for Baracoa River (n = 31). The result for *K* = 2 after Evanno’s method is presented. Individuals identified as *G. punctata* by mtDNA are labeled with arrows and hybrids from Papaya River by dots in the top of the graphic. (B) Analysis of each group independently. In each case the number *K* tested varies and the *K* selected after Evanno’s and these showing relevant biological clusters above the Δ*K* value are presented.

### 3.4 Genetic structuration of populations within groups

Within groups, mtDNA haplotype distribution appeared geographically structured, some haplotypes been not shared by some regions or localities (Fig. 2C). Indeed, most local populations showed unique haplotypes, some haplotypes been shared between neighboring localities (Supplementary material 1a). We also analyzed the microsatellite loci using STRUCTURE to check whether both types of genetic markers reveal similar spatial structures. First, we performed exploratory runs with *K* = 1 to 8 and five repetitions to optimize computation effort. After determining the optimal value of *K* using the program STRUCTURE HARVESTER, 20 replicates were performed.

In *Gambusia rhizophorae*, high *cyt*b distances were found between Mil Cumbres (4) and the other localities (14 mutations between the nearest haplotypes, *p* = 2.5% ± 0.5). In addition, a strong population subdivision was evidenced using STRUCTURE analysis (with *K* = 1 to 8) of microsatellite loci (Fig. 3B; Supplementary material 5b). According to Evanno’s method, *K* = 6 showed the highest probability and distinguished five populations in this group, whose geographic range correspond to single rivers: Camarones (2), San Claudio (5), Baracoa (8, 9), Guanabo (10), Boca de Jaruco (11) and Jibacoa (12). In only one case, El Vajazal Lagoon (1), it was detected mixed membership probabilities with Camarones and Baracoa rivers indicating some shared ancestries.

In *G. punctata*, two major haplogroups (separated by 8 mutations, *p* = 1.6%) were identified, one distributed along south Pinar del Río and another one distributed in La Havana, Matanzas, Isla de la Juventud and La Grifa Key (23) in La Broa Cove. Curiously, haplotypes of both haplogroups were detected in La Siguanea population (26), Isla de la Juventud, suggesting eventual dispersal from western locations or ancestral distribution area (Fig. 2C, Supplementary material 1a). AMOVA analysis detected additional genetic differentiation (Table 2) that is highly congruent with microsatellite results (Fig. 3B). The highest component of variance (73.5%) accounted for differences among geographic areas while 6.4% explained differences among populations inside areas and 20.1% was due to differences inside populations. Four clusters were distinguished; one including the westernmost sampling sites La Papaya and El Convento rivers (13 and 14 respectively) and La Siguanea (26), Isla de la Juventud. The second cluster included localities distributed southeastern Guaniguanico Mountain Range: Los Palacios, San Cristóbal, Masón Stream and San Juan River (15 – 18 in that order). The third one comprised Isla de la Juventud (23, 25, and 26) and La Grifa Key (23). A fourth cluster included all localities from western La Habana to Matanzas provinces (19 – 33). Microsatellite loci analysis, conducted for *K* = 1 to 5 (Supplementary material 5b), distinguished San Juan River (18) population (Evanno’s *K* = 2) from all other localities while a cluster including La Papaya (13) and Itabo (Isla de la Juventud 24) rivers was additionally revealed for *K* = 3. Guanimar Lagoon (19) showed a mixed pattern of individual assignments and the rest of eastern localities (29 – 32) clustered together. For *Gambusia* sp. D, too few mtDNA sequences were available for testing for population structure, however, microsatellite loci analysis conducted with *K* = 1 to 5 (Supplementary material 5b) distinguished two clusters apparently associated with plains and mountain ecosystems. Sampling localities Ariamo River (35), Batata Stream (40) and El Caburní fall (42), which clustered together, are located in highlands while the other localities are in lowlands. Finally, we also considered the two haplotype partitions recognized for *Gambusia* sp. in the eastern region and in the hybrid zone. The first one was geographically delimited by El Brazo River (59) in Camagüey province and included all eastern populations and the second one, was distributed from Calabazas River (44), Sancti Spiritus to Santa María River (58), Camagüey. AMOVA analysis revealed five subgroups, two inside the eastern region and three within the hybrid zone (Table 2). The highest variance component (57.4%, p < 0.0001) explained differences among geographic areas followed by differences among populations (23.7%) and differences inside populations (19%). The hybrid zone showed three clusters: La Jenifer sinkhole (53) in Coco Key (North); Algodón Grande and Cuervo keys (55, 56) in Ana María Gulf; and the rest of the localities of this group (44 - 52, 54, 57 – 58), excluding El Brazo River (59). The easter region showed a cluster containing El Brazo River and the two localities sampled in Yara River (62, 64), and a second cluster including Jobabo, Cerro Colorado, Bayamo and Cojimar (60-61, 63, 65) rivers. Cuabas River (66) was not included in the analysis because the haplotype (R1) found in this locality (n = 2) was common in the hybrid zone but absent from other localities of the eastern region. The STRUCTURE analysis of microsatellite loci applied to the easter region and conducted for *K* = 1 to 5 (Supplementary material 5b), concurred with mtDNA results in revealing three clusters (Fig. 3B). The first cluster was formed by Cojimar River (65) individuals (with *K* = 2 or 3); the second one, containing Yara River individuals and the third one, grouping Cerro Colorado and Bayamo rivers were revealed with *K* = 3. In this case, the two individuals from Cuabas River (66) showed membership probabilities to two different clusters (Fig. 3B).

**Table 2.** The result of AMOVAs using mtDNA sequences to test major geographic haplogroups partitions.

| Source of variation | d.f. | Sum of squares | Variance components | % of variation | Fixation Indices | Significance |
| --- | --- | --- | --- | --- | --- | --- |
| *Gambusia punctata* |  |  |  |  |  |  |
| Among groups | 3 | 116.501 | 4.13029 Va | 73.55 | $F_{CT} = 0.7355$ | $p < 0.044$ |
| Among populations within groups | 2 | 7.302 | 0.35762 Vb | 6.37 | $F_{SC} = 0.2408$ | $p < 0.013$ |
| Inside populations | 35 | 39.465 | 1.12758 Vc | 20.08 | $F_{ST} = 0.7992$ | $p < 0.000$ |
| Total | 40 | 163.268 | 5.6155 |  |  |  |
| *Gambusia* sp. /hybrid zone |  |  |  |  |  |  |
| Among groups | 4 | 45.539 | 1.57853 Va | 68.07 | $F_{CT} = 0.6807$ | $p < 0.000$ |
| Among populations within groups | 15 | 15.393 | 0.24409 Vb | 10.53 | $F_{SC} = 0.3296$ | $p < 0.002$ |
| Inside populations | 23 | 11.417 | 0.49638 Vc | 21.40 | $F_{ST} = 0.7859$ | $p < 0.000$ |
| Total | 42 | 72.349 | 2.3190 |  |  |  |

### 3.5 Genetic diversity and demography

Estimates of genetic diversity are shown in Table 3. Most of the diversity estimates (π, Ho, He, the percentage of polymorphic loci, number of alleles and number of private alleles), as well as the number of differences among haplotypes, were higher in the western groups. *Gambusia rhizophorae* showed a higher number of alleles (9.2, *t*_(96)_ = 4.67, p = 0.00001) and a much higher number of private alleles (n = 15) than *G. punctata* (n = 5), while *Gambusia* sp. D showed the lowest number of private alleles (n = 2) and *Ho* (0.225). *Gambusia punctata* showed higher values of π (0.0109, t_(70)_ = 4.33, p = 0.0005) and mean number of haplotype differences (8.163, t_(70)_ = 3.98, p = 0.0002) than *G. rhizophorae*. The distribution of private alleles was almost evenly among the analyzed localities of *G. rhizophorae* (8/8), less so in *G. punctata* (5/8) and highly skewed on the other two groups: *Gambusia* sp. D (1/6) *Gambusia* sp. (2/5) and hybrid zone (2/6). High and statistically significant *F*_IS_ statistic estimates were obtained for all groups due to lower than expected *H*o values. This clearly reflects a Wahlund effect consequence of the strong population structure observed.

**Table 3:** Diversity estimates obtained for each of the groups. mtDNA estimates for *Gambusia* sp. and Hybrid zone were computed together.

| Group/<br>Statistics | *G. rhizophorae* | *G. punctata* | *Gambusia* sp. D | Hybrid zone | *Gambusia* sp. |
| --- | --- | --- | --- | --- | --- |
| **mtDNA (*cytb*)** |  |  |  |  |  |
| N | 31 | 41 | 21 |  | 44 |
| S | 34 | 39 | 21 |  | 23 |
| H | 15 | 24 | 15 |  | 22 |
| No. differences | 21 | 20 | 9 |  | 11 |
| $K \pm SD$ | 6.955<br>$\pm 1.358$ | 8.163<br>$\pm 1.159$ | 3.943<br>$\pm 1.083$ | | 3.382<br>$\pm 0.910$ |
| $Hd \pm SD$ | 0.923<br>$\pm 0.028$ | 0.880<br>$\pm 0.047$ | 0.957<br>$\pm 0.030$ | | 0.875<br>$\pm 0.044$ |
| $\pi \pm SD$ | 0.0093<br>$\pm 0.0019$ | 0.0109<br>$\pm 0.0009$ | 0.0052<br>$\pm 0.0006$ | | 0.0045<br>$\pm 0.0005$ |
| $F_s$ | - 1.546 ns | - 6.398 * | - 7.715 ** | | - 12.753 *** |
| $D$ | - 0.749 ns | - 0.444 ns | - 0.7881 ns | | - 1.454 ns |
| **Microsatellites** |  |  |  |  |  |
| N | 50 | 48 | 35 | 41 | 38 |
| % Polymorphic loci | 100 | 100 | 88.9 | 77.79 | 88.9 |
| No. Alleles ± SE | 9.2 ± 1.58 | 7.9 ± 1.15 | 4.4 ± 0.96 | 7.2 ± 1.96 | 7.2 ± 2.12 |
| No. Private alleles | 15 | 5 | 2 | 3 | 3 |
| $H_o \pm SE$ | 0.413 ± 0.054 | 0.444 ± 0.070 | 0.225 ± 0.050 | 0.319 ± 0.081 | 0.319 ± 0.078 |
| $He \pm SE$ | 0.702 ± 0.075 | 0.634 ± 0.081 | 0.440 ± 0.101 | 0.465 ± 0.114 | 0.499 ± 0.117 |
| $F_{IS}$ | 0.382 ± 0.067* | 0.280 ± 0.059* | 0.379 ± 0.092* | 0.318 ± 0.047* | 0.310 ± 0.064* |

N is the sample size; S is the number of segregating sites; h is the number of haplotypes; No. differences is the maximum number of differences observed among haplotypes; $K$ is the mean number of differences among haplotypes; $Hd$ is the haplotype diversity; $\pi$ is the nucleotide diversity (Nei, 1987); $F_s$ is Fu's test (Fu, 1997); $D$ is Tajima's $D$ statistic (Tajima, 1989); $H_o$ and $He$ are the observed and expected heterozygosity and $F_{IS}$ is the fixation index according to Weir and Cockerham (1984). \*( $p < 0.05$ ); \*\* ( $p < 0.01$ ); \*\*\* ( $p < 0.001$ ) obtained after 10,000 coalescence simulations.

We used Tajima’s *D* and Fu’s *F*_S_ neutrality test to detect past population expansions on the different groups. In all cases, Tajima’s *D* tests were negative and failed to detect any departure from expected proportions under neutrality assumptions. However, *F*_S_ test was statistically significant in all but *G. rhizophorae* group showing an eastward increase of magnitude (- 1.546 in *G. rhizophorae* to - 12.753 in *Gambusia* sp. (Table 3). This pattern suggests that the *cyt*b sequences are evolving under neutrality and that population expansions might have occurred in three of the groups.

## 4. Discussion

With this study, we provide further evidence indicating that genetic differentiation in *Gambusia punctata* species group in Cuba is higher than expected using morphological characters alone (Rivas, 1969). Moreover, these genetically differentiated groups are well separated geographically and they are currently occupying areas corresponding to the major ancient land cores of the Archipelago and their current-time dryland surroundings.

### 4.1 Cryptic diversity

The lack of obvious morphological differentiation among cryptic species had often hampered an accurate estimation of the biodiversity and biased our understanding of some evolutionary processes (Fišer et al., 2018). Indeed, molecular markers have evidenced that hidden whereas structured genetic diversity is common in most animal phyla (Bickford et al., 2007). Recently, different studies have pointed out that the genetic diversity of the Cuban freshwater fishes have been largely underestimated and that new candidate species or subspecies actually occurs in genera such as *Girardinus* (Doadrio et al., 2009), *Gambusia* (Lara et al., 2010), *Lucifuga* (García-Machado et al., 2011), *Rivulus* (Ponce de León et al., 2014; Rodríguez, 2015b), *Dormitator* (Galván-Quesada et al., 2016) and *Kryptolebias* (Tatarenkov et al., 2017).

Compared to other poeciliid genera, *Gambusia* species diversification has been relatively less pronounced in the Caribbean islands (Lucinda, 2003). Whereas *Limia* in Hispaniola (Weaver et al., 2016) and *Girardinus* in Cuba (Doadrio et al., 2009; Lara et al., 2010; Rivas, 1958) display impressive radiations, *Gambusia* species groups (i.e. *punctata* and *puncticulata*) show a relatively larger geographic distribution and a lower number of species per island (Rauchenberger, 1989). This pattern could result from higher dispersal capabilities and/or more recent colonization of the Caribbean by *Gambusia* compared to the other poeciliid genera (Reznick et al., 2017) but it may just reflect the lack of obvious morphological diversity in these groups. Here we provide evidence for new putative species inside *G. punctata* group in Cuba supporting the idea that the diversity of the group has been underestimated. Integrating the results of mtDNA and nuclear microsatellite genetic markers and geographic distributions we show that the species *G. punctata* in Cuba is subdivided into four independent cryptic entities showing parapatric distributions. In addition to the nominal *G. punctata* and *G. rhizophorae*, both found in western regions of the island, we identified three new lineages. *Gambusia* sp., first evidenced as a well-differentiated mtDNA haplogroup (Lara et al., 2010) from east-central Cuba, was also supported by the microsatellite loci analysis in the present study. However, whereas Lara et al. (2010) included samples from Coco Key as part of this group, we delimited the geographic distribution of *Gambusia* sp. to the eastern (i.e. Holguin – Santiago de Cuba) region. The second group, *Gambusia* sp. D, distributed into Guamuhaya mountain range and nearby areas in the central region of the island, was supported by both mitochondrial and microsatellite data. The area in between central and eastern regions represents a large area of genetic intergradations and hybridization between these two entities (the hybrid zone, see below). Finally, we found evidence for a fifth putative cryptic species, restricted to a single locality at the mountainous area of Mil Cumbres, and which represents a divergent mtDNA subclade within *G. rhizophorae*. The mtDNA phylogenetic analysis suggests that *Gambusia* sp. D and *Gambusia* sp. are not sister groups. However, even if they are they are distantly related (see Figure 2 and Supplementary material 2). We should note that sequences from multiple independent nuclear genes are necessary in order to test the phylogenetic hypothesis proposed here with the mtDNA sequences.

The PTP model analysis based on the mtDNA sequences provided strong support for *Gambusia* sp. while the other clades were not. However, this delimitation can only be considered as a preliminary hypothesis of species that need corroboration (Zhang et al., 2013). Lumping of clades (Kozak et al., 2015) or a tendency to overestimate the true number of species (Postaire et al., 2016), have been observed when using species delimitation methods. Nonetheless, the five groups can be unambiguously diagnosed by a combination of 44 nucleotide sites and private DNA positions at *cyt*b (Supplementary material 6). Similarly, the four lineages (Mil Cumbres individuals not included in the analysis) were also strongly supported by the results of the microsatellite *loci* analysis with the program STRUCTURE. This analysis showed homogeneous clusters of individuals showing very little or no admixture between the groups out of the contact zones.

Genetic distances, using *cyt*b or COI partial sequences, were in all cases higher than 2% (2.7 – 6.9%, see Table 1), figures that suggest species-level divergence in most fishes (April et al., 2011; Johns and Avise, 1998; Pereira et al., 2013). In the same way, Jost’s D statistic estimates based on microsatellite loci were high in all group pairwise comparisons and *F*_ST_ values were in the same range than those estimated between *Gambusia affinis* and *G. holbrooki*, two species that currently hybridize in south-east North America (Wilk and Horth, 2016) and that show a *cyt*b distance around 4.0% (based on *p* distances between 28 *G. holbrooki* and 9 *G. affinis* 394 bp sequences retrieved from GenBank). In order to distinguish intraspecific and interspecific levels of divergence, we have extensively sampled along the distribution area of the nominal species *G. punctata* and *G. rhizophorae*. This strategy revealed that each species is indeed an arrangement of populations showing, in some cases, strong geographic structures. However, intra-species genetic distances were lower than between species.

Although we did not reevaluate morphological characters, in a previous study Rivas (1969) analyzed a large number of individuals of *G. punctata* (n = 1319) from 37 localities distributed along Cuba and Isla de La Juventud. Rivas found no evidence of morphological partitions inside this species but a substantial overlap of morphological variation between *G. punctata* and *G. rhizophorae* has found. Recently, Fišer et al. (2018) reviewed three possible mechanisms that have been proposed as generators of cryptic diversity: recent divergence, morphological convergence, and niche conservatism. In order to test their specific predictions and hence to disentangle their role in the evolution of the *G. punctata* species group, accurate estimates of species ages as well as information about their ecology and the analysis of the possible selective factors intervening during the evolution of the group are mandatory. With the available data, we speculate that a recent divergence might not be the main mechanism. Using a rough approximation of nucleotide substitutions of 1% per million years at the *cyt*b in fishes (Bermingham et al., 1997) these groups have diverged between 7 and 4 Mya. Similar estimates (10.4 – 4.4 Mya) were obtained between *G. punctata* and *G. rhizophorae* in a previous study using several nuclear and mitochondrial gene sequences (Reznick et al., 2017). This implies a large time-lapse for accumulating morphological differences. The generalist ecological niches described for *Gambusia punctata* species group in Cuba *sensu* Rivas 1969 (Ponce de León and Rodríguez, 2010; Ponce de León and Rodríguez, 2013; Rodríguez-Machado et al., 2019) point to niche conservatism as a mechanism that could be involve in morphological stasis.

### 4.2 Salinity as a driver of speciation

The geological history of the Cuban archipelago brings a remarkably attractive framework to understand *Gambusia* fish biogeography in the Caribbean. As the land cores that represented the Cuban palaeoarchipelago evolved disconnected to Central and North America, it implies that the *Gambusia* species colonizing these areas were salt-tolerant, at least temporarily. Salt tolerance is widely recognized in poeciliids, and particularly in *Gambusia* species, as a factor explaining several possible cases of dispersal throughout marine barriers including the colonization of Cuban archipelago (Myers, 1938; Ponce de León et al., 2014; Reznick et al., 2017; Rosen and Bailey, 1963). Our results suggest that all Cuban members of the *Gambusia punctata* species group are salt-tolerant, contrary to the hypothesis proposed by Rivas (1969). Adaptation to different ecological conditions, such as different level of salinity, has been recognized as important factors promoting divergence of freshwater fish populations (Langerhans et al., 2007; McGee et al., 2013; Perreault-Payette et al., 2017; Tobler et al., 2018; Vines and Schluter, 2006), and ultimately speciation such as in the Baltic cod (Berg et al., 2015) and European flounders (Momigliano et al., 2017). Early evidence that *G. punctata* and *G. rhizophorae* live in allopatry (and parapatry at some localities) and are adapted to different environments (Rivas, 1969), suggested ecological speciation due to the adaptation to different salinity levels, and reproductive isolation as a by-product. However, in all major clades of the *Gambusia punctata* species group found in Cuba (i.e. *Gambusia punctata*, *G. rhizophorae*, *Gambusia* sp. D and *Gambusia* sp.), we collected individuals inhabiting at least two of the three - freshwater, brackish or saltwater - habitats (Supplementary material 1). This is consistent with previous findings such as the presence of two *Gambusia* sp. populations inhabiting waters with contrasting salinity levels (Lara et al., 2010). All these findings suggest that adaptation to different salinity levels had not been a key factor involved in the geographic differentiation of putative species in the *Gambusia punctata* species group in Cuba. More generally, rather than ecological speciation due to the adaptation to different ecological conditions, speciation in the *Gambusia punctata* species group in Cuba seems more likely determined by long periods of isolation due to geological and climatic events shaping the topography of the archipelago and the connectivity of land habitats.

### 4.3 Phylogeny and phylogeographic considerations

The phylogenies obtained in the present study suggest that the species group diversification in Cuba might have occurred in a relatively small period of time, as indicated by the short length of the branch subtending the earliest divergences within the group, followed by a relatively long period of time of allopatric isolation. The current geographic distribution of the groups largely matches the emerged land cores composing the proto archipelago during upper Miocene and Late Pliocene (Iturralde-Vinent, 2006; Iturralde-Vinent and MacPhee, 1999), and it is consistent with the period estimated (10.4 – 4.4 Ma) for the split of *G. rhizophorae* and *G. punctata* proposed by Reznick et al. (2017). Although these dating should be considered only tentative, they imply that the ancestor was in Cuba by later Miocene - early Pliocene epochs. This suggests that the diversification within the *Gambusia punctata* species group in Cuba may have occurred, at least to some extent, related to the isolation of land cores caused by geological events and fluctuations of the sea level during the late Miocene and early Pliocene (Iturralde-Vinent, 2006; Iturralde-Vinent and MacPhee, 1999). Unexpectedly, salinity tolerance did not seem to have promoted homogenization of populations at the scale of the Cuban archipelago.

Similar geographic distributions have been found in other animals such as frogs (Rodríguez et al., 2013; Rodríguez et al., 2010), toads (Alonso et al., 2012), anoles (Glor et al., 2004), and plants such as *Leucocroton* (Jestrow et al., 2012). Interestingly, the geographic distributions of *Gambusia punctata* and *G. rhizophorae* match with those of *Rivulus cylindraceus* and *R. berovidesi* (Ponce de León et al., 2014; Rodríguez, 2015b), and the phylogeographic distribution of mtDNA haplotypes of *G. punctata* and *R. cylindraceus* are very similar (Ponce de León et al., 2014). The finding that the haplotypes distinguishing western and eastern haplogroups in *G. puntata* co-occur in La Siguanea (Isla de la Juventud), raised the hypothesis that *G. punctata* and *G. rhizophorae* have diverged in allopatry, the first within Isla de la Juventud and has dispersed later to the main island as in the case of *R. cylindraceus* (Ponce de León et al., 2014). These examples pointed to common evolutionary histories involving the geological evolution of the Cuban archipelago.

Although a deep analysis of the demographic history of the populations is out of the scope of the present study, preliminary inferences provided some clues about the recent past demographic history of the different groups. *F*s tests suggest population growth for three of the groups: *G. punctata*, *Gambusia* sp. D and *Gambusia* sp. This pattern can be related to recurrent periods of land habitat contractions or loss as a result of marine transgressions or desiccation of lakes and rivers due to low precipitations in the Caribbean region during the last ten thousand years (Hodell et al., 2000). The patterns of haplotype and nucleotide diversity could also reflect expansion in some cases. A combination of high *H*d and low π observed in *Gambusia* sp. D and *Gambusia* sp., which showed the highest *F*s values, is typical of a rapid population expansion after a bottleneck (Grant and Bowen, 1998). This might have strengthened population isolation, as observed within each group, as well as allopatric divergence between the groups. However, as the neutrality test used here can be confounded by selection and population fragmentation these results should be considered cautiously. For instance, directional selection and weak purifying selection may mimic the effects of population expansion (Ballard and Dean, 2001; Braverman et al., 1995). Similarly, population subdivision can upward bias in the number of singleton variants in haploid genomes (Hammer et al., 2003).

Another remarkable result of the present study is the finding that the available sequences of *G. rhizophorae* from Florida are phylogenetically nested within the Cuban clade of this species, which implies that *G. rhizophorae* colonized Florida after a relatively recent over-water dispersal event from the island to the mainland. This challenge the hypothesis suggesting a North American ancestor of the group (Reznick et al., 2017). Given the high tolerance to saltwater, the patterns of marine currents and the high frequency of meteorological phenomena like hurricanes and storms, it is to some extent expected that this type of dispersal events happened repeatedly and thus, might have facilitated the dispersion of *Gambusia* in the Caribbean area. Such dispersal events could have also occurred between the main island and Isla de la Juventud in *G. punctata* (haplotype B14 shared between Papaya and Convento rivers and La Siguanea) and *Rivulus* (Ponce de León et al., 2014), and between the hybrid zone and nearby keys (haplotype R13 shared between Tayabacoa and Majagua rivers and Algodón Grande Key). Although we cannot overrule the possibility of man-mediated dispersal events, these examples strengthens the view that the Greater Antillean islands may represent centers of origin for regional species diversity (Glor et al., 2005) and integrates the list of cases supporting colonization of mainland by animal and plant species from Caribbean island ancestors as a result of overseas dispersal events (Glor et al., 2005; Gugger and Cavender-Bares, 2013; Heinicke et al., 2011; Nicholson et al., 2005).

In spite of the high dispersal abilities of these species, we revealed strong genetic structures between groups and within each group suggesting that the interplay between historical and local factors might have shaped the connectivity among areas at different scales producing isolation and reconnection across time. In addition, most of the streams and river’s watercourses in Cuba flows straight towards the sea and shows few interconnections between one and another, favoring local isolation of fish populations.

### 4.4 Cases of hybridization and introgression

Hybridization has long been perceived as evidence of the absence of intrinsic reproductive isolation between populations and, in consequence, as an indicator of lack of independent evolution (Dobzhansky, 1950; Mayr, 1942). However, it is increasingly clear that this process is widespread between groups rapidly radiating (Mallet, 2007). Our results have evidenced that hybridization is part of the evolutionary processes operating in the *G. punctata* species group in Cuba. Three cases of hybridization were detected; two between *G. punctata* and *G. rhizophorae*, restricted to single localities and the third between *Gambusia* sp. D and *Gambusia* sp. occupying a larger area of the east-central region. Considering the current distributions of these groups, the scenario suggested involves secondary contacts after range expansion and incomplete reproductive barriers (Carson and Dowling, 2006; Petit and Excoffier, 2009). In each case, asymmetric mtDNA introgression was detected suggesting that some reproductive bias or selection have potentially taken place (Meyer et al., 2006). In *G. punctata* and *G. rhizophorae* contact sites, Papaya and Baracoa rivers, the mtDNA of *G. punctata* was present in the population. In Papaya River, all individuals carried *G. punctata* mtDNA while at Baracoa River they represented the 22.6%. However, while all individuals were assigned unequivocally to *G. rhizophorae* at Baracoa River, admixed ancestries were found at Papaya River. This suggests that population interactions might differ in these cases.

In the case of *Gambusia* sp. D and *Gambusia* sp., the hybrid zone is much wider. We have found a complete replacement of *Gambusia* sp. D mtDNA by *Gambusia* sp. mtDNA genotypes all over the area, while we found a mixture of nuclear genotypes with a spatial arrangement correlated with the distance to the parent species. The geographic diversification of the mtDNA hybrid zone and the distinction between parent groups and hybrid mtDNA haplotypes and nuclear loci suggests that the origin of the contact was not recent and that the hybrid zone has existed for some time, a pattern also observed in other fish species (e.g. Carson and Dowling, 2006). The example described here appears similar to that found between *Gambusia affinis* and *G. holbrooki*, two species that hybridize at their stable contact zone in the southern United States (Scribner and Avise, 1993; Wilk and Horth, 2016).

### 4.5 Taxonomic insights

Given the strong genetic differentiation among groups and well-separated geographic distributions, we propose to consider *Gambusia* sp. D and *Gambusia* sp. as putative species in addition to *G. punctata* and *G. rhizophorae*. We provide a set of molecular markers unique to each of the four lineages allowing their identification. Additionally, the cryptic lineage of *G. rhizophorae* from Mil Cumbres although strongly divergent by the mtDNA and well delimited by a unique arrangement of diagnostic nucleotide substitutions need further corroboration by the analysis of additional data (e.g. nuclear genes). Detailed morphological studies and, ideally, formal descriptions of these putative new species are now necessary to put them on the focus of management and conservation authorities.

## 5. Acknowledgments

Authors are sincerely grateful to Martin Acosta, Lourdes Mugica and Fabian Pina for providing logistic support. We thank also the two anonymous reviewers and the editor for their valuable comments and suggestions that have contributed to improving the document. We would also like to thank Norvis Hernández for the help in the field and Fatima Shahid and Aidan Small for reviewing the language.

## References

Alonso, R., Crawford, A., Bermingham, E., 2012. Molecular phylogeny of an endemic radiation of Cuban toads (Bufonidae: Peltophryne) based on mitochondrial and nuclear genes. J. Biogeogr. 39, 434–451. https://doi.org/10.1111/j.1365-2699.2011.02594.x.

April, J., Mayden, R.L., Hanner, R.H., Bernatchez, L., 2011. Genetic calibration of species diversity among North America’s freshwater fishes. PNAS 108, 10602–10607. http://doi.org/10.1073/pnas.1016437108.

Ballard, J.W.O., Dean, M.D., 2001. The mitochondrial genome: mutation, selection and recombination. Current Opinion in Genetics & Development 11, 667–672. https://doi.org/10.1016/S0959-437X(00)00251-3.

Bandelt, H.J., Forster, P., Röhl, A., 1999. Median-joining networks for inferring intraspecific phylogenies. Molecular Biology and Evolution 16, 37–48. http://doi.org/10.1093/oxfordjournals.molbev.a026036.

Berg, P.R., Jentoft, S., Star, B., Ring, K.H., Knutsen, H., Lien, S., Jakobsen, K.S., André, C., 2015. Adaptation to low salinity promotes genomic divergence in Atlantic Cod (*Gadus morhua* L.). Genome Biol. Evol. 7, 1644–1663. http://doi.org/10.1093/gbe/evv093.

Bermingham, E., McCafferty, S., Martin, A., 1997. Fish biogeography and molecular clocks: perspectives from the Panamanian Isthmus. In: Kocher, T., Stepien, C. (Eds.), Molecular Systematics of Fishes. Academic, New York.

Bickford, D., Lohman, D.J., Sodhi, N.S., Ng, P.K.L., Meier, R., Winker, K., Ingram, K.K., Das, I., 2007. Cryptic species as a window on diversity and conservation. TRENDS in Ecology and Evolution 22, 148–155. https://doi.org/10.1016/j.tree.2006.11.004.

Braverman, J., Hudson, R., Kaplan, N., Langley, C., Stephan, W., 1995. The hitchhiking effect on the site frequency spectrum of DNA polymorphisms. Genetics 140, 783–796.

Carson, E.W., Dowling, T.E., 2006. Influence of hydrogeographic history and hybridization on the distribution of genetic variation in the pupfishes *Cyprinodon atrorus* and *C. bifasciatus*. Molecular Ecology 15, 667–679. http://doi.org/10.1111/j.1365-294X.2005.02763.x.

Cureton, J.C., Buchman, A.B., Randle, C., Lutterschmidt, W.I., Deaton, R., 2010. Development and cross-amplification of nine novel *Gambusia geiseri* microsatellite loci in *G. clark-hubbsi* and the endangered *G. nobilis*. Conservation Genet Resour 2, 177–179. https://doi.org/10.1007/s12686-010-9220-7.

Doadrio, I., Perea, S., Alcaraz, L., Hernández, N., 2009. Molecular phylogeny and biogeography of the Cuban genus *Girardinus* Poey, 1854 and relationships within the tribe Girardinini (Actinopterygii, Poeciliidae). Molecular Phylogenetics and Evolution 50, 16–30. http://doi.org/10.1016/j.ympev.2008.09.014.

Dobzhansky, T., 1950. Mendelian populations and their evolution. The American Naturalist 84, 401–418.

Earl, D.A., vonHoldt, B.M., 2012. STRUCTURE HARVESTER: a website and program for visualizing STRUCTURE output and implementing the Evanno method. Conserv. Gen. Resour. 4, 359–361. 10.1007/s12686-011-9548-7.

Evanno, G., Regnaut, S., Goudet, J., 2005. Detecting the number of clusters of individuals using the software STRUCTURE: a simulation study. Molecular Ecology 14, 2611–2620. http://doi.org/10.1111/j.1365-294X.2005.02553.x.

Excoffier, L., Lischer, H.E.L., 2010. Arlequin suite ver 3.5: a new series of programs to perform population genetics analyses under Linux and Windows. Molecular Ecology Resources 10, 564–567. http://doi.org/10.1111/j.1755-0998.2010.02847.x.

Excoffier, L., Smouse, P.E., Quattro, J.M., 1992. Analysis of molecular variance inferred from metric distances among DNA haplotypes: Application to human mitochondrial DNA restriction data. Genetics 131, 479–491.

Fink, W., 1971. A revision of the *Gambusia puncticulata* complex (Pisces: Poecilidae). Publication of the Gulf Coast Research Laboratory Museum 2, 11–46.

Fišer, C., Robinson, C.T., Malard, F., 2018. Cryptic species as a window into the paradigm shift of the species concept. Molecular Ecology 27, 613–635. http://doi.org/10.1111/mec.14486.

Fu, Y.-X., 1997. Statistical tests of neutrality of mutations against population growth, hitchhiking and background selection. Genetics 147, 915–925.

Galván-Quesada, S., Doadrio, I., Alda, F., Perdices, A., Reina, R.G., García Varela, M., Hernández, N., Campos Mendoza, A., Bermingham, E., Domínguez-Domínguez, O., 2016. Molecular phylogeny and biogeography of the amphidromous fish genus *Dormitator* Gill 1861 (Teleostei: Eleotridae). PLOS ONE 11, e0153538. http://doi.org/10.1371/journal.pone.0153538.

García-Machado, E., Hernández, D., García-Debrás, A., Chevalier-Monteagudo, P., Metcalfe, C., Bernatchez, L., Casane, D., 2011. Molecular phylogeny and phylogeography of the Cuban cave-fishes of the genus *Lucifuga*: Evidence for cryptic allopatric diversity. Molecular Phylogenetics and Evolution 61, 470–483. http://doi.org/10.1016/j.ympev.2011.06.015.

Glor, R., Gifford, M., Larson, A., Losos, J., Rodríguez Chetino, L., Chamizo Lara, A., Jackman, T., 2004. Partial island submergence and speciation in an adaptative radiation: a multilocus analysis of the Cuban green anoles. Proc. R. Soc. Lond. B 271, 2257–2265. https://doi.org/10.1098/rspb.2004.2819.

Glor, R., Losos, J., Larson, A., 2005. Out of Cuba: overwater dispersal and speciation among lizards in the *Anolis carolinensis* subgroup. Molecular Ecology 14, 2419–2432. http://doi.org/10.1111/j.1365-294X.2005.02550.x.

Goudet, J., 2001. “FSTAT, a program to estimate and test gene diversities and fixation indices” (version 2.9.3). http://www.unil.ch/izea/softwares/fstat.html.

Grant, W., Bowen, B., 1998. Shallow population histories in deep evolutionary lineages of marine fishes: Insights from sardines and anchovies and lessons for conservation. Journal of Heredity 89 (5), 415–426.

Gugger, P.F., Cavender-Bares, J., 2013. Molecular and morphological support for a Florida origin of the Cuban oak. Journal of Biogeography 40, 632–645. http://doi.org/10.1111/j.1365-2699.2011.02610.x.

Hall, T.A., 1999. BioEdit: a user-friendly biological sequence alignment editor and analysis program for Windows 95/98/NT. Nucleics Acids Symposium Series 41, 95–98.

Hammer, M.F., Blackmer, F., Garrigan, D., Nachman, M.W., Wilder, J.A., 2003. Human population structure and its effects on sampling Y chromosome sequence variation. Genetics 164, 1495–1509.

Heinen-Kay, J.L., Noel, H.G., Layman, C.A., Langerhans, R.B., 2014. Human-caused habitat fragmentation can drive rapid divergence of male genitalia. Evolutionary Applications 7, 1252–1267. http://doi.org/10.1111/eva.12223.

Heinicke, M.P., Diaz, L.M., Hedges, S.B., 2011. Origin of invasive Florida frogs traced to Cuba. Biology letters 7, 407–410. http://doi.org/10.1098/rsbl.2010.1131.

Hernández, D., Casane, D., Chevalier-Monteagudo, P., Bernatchez, L., García-Machado, E., 2016. Go West: A One Way Stepping-Stone Dispersion Model for the Cavefish Lucifuga dentata in Western Cuba. PLOS ONE 11, e0153545. 10.1371/journal.pone.0153545.

Hodell, D.A., Brenner, M., Curtis, J.H., 2000. Climate Changes in the Northern American Tropics and Subtropics since the Last Ice Age: Implications for Environment and Culture. In: Lents, D.L. (Ed.), Imperfect Balance: Landscape Transformations in the Precolumbian Americas. Columbia University Press, New York., pp. 13–38

Iturralde-Vinent, M., 2006. Meso-Cenozoic Caribbean paleogeography: Implications for the historical biogeography of the region. International Geology Review 48, 791–827. http://doi.org/10.2747/0020-6814.48.9.791.

Iturralde-Vinent, M., MacPhee, R.D.E., 1999. Paleogeography of the Caribbean region: Implications for Cenozoic biogeography. American Museum of Natural History Bulletin 238, 1–95.

Jakobsson, M., Rosenberg, N.A., 2007. CLUMPP: a cluster matching and permutation program for dealing with label switching and multimodality in analysis of population structure. Bioinformatics 23, 1801–1806. http://doi.org/10.1093/bioinformatics/btm233.

Jestrow, B., Gutiérrez Amaro, J., Francisco-Ortega, J., 2012. Islands within islands: a molecular phylogenetic study of the *Leucocroton alliance* (Euphorbiaceae) across the Caribbean Islands and within the serpentinite archipelago of Cuba. Journal of Biogeography 39, 452–464. http://doi.org/10.1111/j.1365-2699.2011.02607.x.

Johns, G.C., Avise, J.C., 1998. A comparative summary of genetic distances in the vertebrates from the mitochondrial cytochrome b gene. Molecular Biology and Evolution 15, 1481–1490.

Jost, L., 2008. GST and its relatives do not measure differentiation. Molecular Ecology 17, 4015–4026. http://doi.org/S10.1111/j.1365-294X.2008.03887.x.

Jost, L., 2009. D vs. GST: Response to Heller and Siegismund (2009) and Ryman and Leimar (2009). Molecular Ecology 18, 2088–2091. http://doi.org/10.1111/j.1365-294X.2009.04186.x.

Jost, L., Archer, F., Flanagan, S., Gaggiotti, O., Hoban, S., Latch, E., 2018. Differentiation measures for conservation genetics. Evolutionary Applications 11, 1139–1148. http://doi.org/10.1111/eva.12590.

Kozak, K.M., Wahlberg, N., Neild, A.F.E., Dasmahapatra, K.K., Mallet, J., Jiggins, C.D., 2015. Multilocus Species Trees Show the Recent Adaptive Radiation of the Mimetic Heliconius Butterflies. Systematic Biology 64, 505–524. 10.1093/sysbio/syv007.

Kumar, S., Stecher, G., Tamura, K., 2016. MEGA7: Molecular Evolutionary Genetics Analysis Version 7.0 for Bigger Datasets. Molecular Biology and Evolution 33, 1870–1874. http://doi.org/10.1093/molbev/msw054.

Langerhans, R., Gifford, M., Joseph, E., 2007. Ecological speciation in *Gambusia* fishes. Evolution 61-9, 2056–2074. https://doi.org/10.1111/j.1558-5646.2007.00171.x.

Lara, A., Ponce de León, J.L., Rodríguez, R., Casane, D., Côté, G., Bernatchez, L., García-Machado, E., 2010. DNA barcoding of Cuban freshwater fishes: evidence for cryptic species and taxonomic conflicts. Molecular Ecology Resources 10, 421–430. http://doi.org/10.1111/j.1755-0998.2009.02785.x.

Librado, P., Rozas, J., 2009. DnaSP v5: A software for comprehensive analysis of DNA polymorphism data. BIOINFORMATICS 25, 1451–1452.

Lucinda, P.H.F., 2003. Family Poeciliidae. In: Reis, R.E., Kullander, S.O., Ferraris, C.J. (Eds.), Check List of the Freshwater Fishes. Edipucrs, 729p, Porto Alegre, pp. 555–581.

Lydeard, C., M.C. Wooten, A. Meyer, 1995. Cytochrome b sequence variation and a molecular phylogeny of the live-beering fish genus *Gambusia* (Cyprinodontiformes: Poeciliidae). Canadian Journal of Zoology 73, 213–227.

Mallet, J., 2007. Hybrid speciation. Nature 446, 279–283. http://doi.org/10.1038/nature05706.

Mayr, E., 1942. Systematics and the origin of species, from the viewpoint of a zoologist. Harvard University Press, Cambridge, MA.

McGee, M.D., Schluter, D., Wainwright, P.C., 2013. Functional basis of ecological divergence in sympatric stickleback. BMC Evolutionary Biology 13, 227. http://doi.org/10.1186.1186/1471-2148-13-277.

Meirmans, P.G., Hedrick, P.W., 2011. Assessing population structure: FST and related measures. Molecular Ecology Resources 11, 5–18. http://doi.org/10.1111/j.1755-0998.2010.02927.x.

Meirmans, P.G., Van Tienderen, P.H., 2004. Genotype and genodive: two programs for the analysis of genetic diversity of asexual organisms. Molecular Ecology Notes 4, 792–794. http://doi.org/10.1111/j.1471-8286.2004.00770.x.

Meyer, A., Kocher, T.D., Basasibwaki, P., Wilson, A.C., 1990. Monophyletic origin of Lake Victoria cichlid fishes suggested by mitochondrial DNA sequences. Nature 374, 550–553.

Meyer, A., Salzburger, W., Schartl, M., 2006. Hybrid origin of a swordtail species (Teleostei: *Xiphophorus clemenciae*) driven by sexual selection. Molecular Ecology 15, 721–730. http://doi.org/10.1111/j.1365-294X.2006.02810.x.

Momigliano, P., Jokinen, H., Fraimout, A., Florin, A.-B., Norkko, A., Merilä, J., 2017. Extraordinarily rapid speciation in a marine fish. PNAS 114, 6074–6079. http://doi.org/10.1073/pnas.1615109114.

Moritz, C., Cicero, C., 2004. DNA Barcoding: Promise and Pitfalls. PLOS Biology 2, e354. 10.1371/journal.pbio.0020354.

Myers, G., 1938. Fresh-water fishes and West Indian zoogeography. Annual Report of the Board of Regents of the Smithsonian Institution 92, 339–364.

Nei, M., 1987. Molecular Evolutionary Genetics. Columbia University Press, New York.

Nicholson, K.E., Glor, R.E., Kolbe, J.J., Larson, A., Blair Hedges, S., Losos, J.B., 2005. Mainland colonization by island lizards. J. Biogeogr. 32, 929–938. http://doi.org/10.1111/j.1365-2699.2004.01222.x.

Palumbi, S.P., 1996. Nucleic acids II: the polymerase chain reaction. In: Hillis, D.M., Moritz, C., Mable, B.K. (Eds.), Molecular Systematics. Sunderland, MA: Sinauer Associates, pp. 205–247.

Pereira, L.H.G., Hanner, R., Foresti, F., Oliveira, C., 2013. Can DNA barcoding accurately discriminate megadiverse Neotropical freshwater fish fauna? BMC Genetics 14, 20–20. http://doi.org/10.1186/1471-2156-14-20.

Perreault-Payette, A., Muir, A.M., Goetz, F., Perrier, C., Normandeau, E., Sirois, P., Bernatchez, L., 2017. Investigating the extent of parallelism in morphological and genomic divergence among lake trout ecotypes in Lake Superior. Molecular Ecology 26, 1477–1497. http://doi.org/10.1111/mec.14018.

Petit, R.J., Excoffier, L., 2009. Gene flow and species delimitation. TRENDS in Ecology and Evolution 24, 386–393. http://doi.org/10.1016/j.tree.2009.02.011.

Polzin, T., Daneshmand, S.V., 2003. On Steiner trees and minimum spanning trees in hypergraphs. Operations Research Letters 31, 12–20. https://doi.org/10.1016/S0167-6377(02)00185-2.

Ponce de León, J., León-Finalé, G., Rodríguez, R., Metcalfe, C., Hernández, D., Casane, D., García-Machado, E., 2014. Phylogeography of Cuban *Rivulus*: evidence for allopatric speciation and secondary dispersal across a marine barrier. Molecular Phylogenetics and Evolution 79 404–414. http://doi.org/10.1016/j.ympev.2014.07.007.

Ponce de León, J., Rodríguez, R., 2010. Peces cubanos de la familia Poeciliidae. Guía de Campo. Editorial Academia, La Habana.

Ponce de León, J.L., Rodríguez, R., 2013. Spatial segregation of freshwater fish in an intermittent Cuban stream. Revista Biología 22, 31–35.

Postaire, B., Magalon, H., Bourmaud, C.A.F., Bruggemann, J.H., 2016. Molecular species delimitation methods and population genetics data reveal extensive lineage diversity and cryptic species in Aglaopheniidae (Hydrozoa). Molecular Phylogenetics and Evolution 105, 36–49. https://doi.org/10.1016/j.ympev.2016.08.013.

Pritchard, J.K., Stephens, M., Donnelly, P., 2000. Inference of population structure using multilocus genotype data. Genetics 155, 945–959.

Purcell, K.M., Lance, S.L., Jones, K.L., Stockwell, C.A., 2011. Ten novel microsatellite markers for the western mosquitofish *Gambusia affinis*. Conservation Genet Resour 3, 361–663. https://doi.org/10.1007/s12686-010-9362-7.

Rauchenberger, M., 1989. Systematics and Biogeography of the genus *Gambusia* (Cyprinodontiformes: Poeciliidae). American Museum Novitates 2951, 1–71.

Reznick, D.N., Furness, A.I., Meredith, R.W., Springer, M.S., 2017. The origin and biogeographic diversification of fishes in the family Poeciliidae. PLOS ONE 12, e0172546. http://doi.org/10.1371/journal.pone.0172546.

Rivas, L.R., 1958. The origin, evolution, dispersal, and geographical distribution of the Cuban poeciliid fishes of the Tribe Girardinini. Proceedings of the American Philosophical Society 102, 281–320.

Rivas, L.R., 1969. A revision of the poeciliid fishes of the *Gambusia punctata* species group with descriptions of two new species. Copeia 1969, 778–795.

Rodríguez-Machado, S., Hidalgo, G., Brito, L., Arrebola, M., Morales, L., Hernández, D., Sánchez, D., Rodríguez-González, J., Castellini, C., Ponce de León, J., 2019. Partial trophic segregation in co-occurring Gambusia species (Cyprinodontiformes: Poeciliidae) in a natural wetland of Cuba. Revista Investigaciones Marinas 39, 76–89.

Rodríguez-Machado, S., Ponce de León, J.L., Casane, D., García-Machado, E., 2017. Morphology and genetics reveal the occurrence of *Girardinus falcatus* (Eigenmann, 1903) (Cyprinodontiformes, Poeciliidae) in eastern Cuba. Check List 13, 1059–1065.

Rodríguez, A., Poth, D., Schulz, S., Gehara, M., Vences, M., 2013. Genetic diversity, phylogeny and evolution of alkaloid sequestering in Cuban miniaturized frogs of the *Eleutherodactylus limbatus* group. Molecular Phylogenetics and Evolution 68, 541–554. https://doi.org/10.1016/j.ympev.2013.04.031.

Rodríguez, A., Vences, M., Nevado, B., Machordom, A., Verheyen, E., 2010. Biogeographic origin and radiation of Cuban *Eleutherodactylus* frogs of the auriculatus species group, inferred from mitochondrial and nuclear gene sequences. Molecular Phylogenetics and Evolution 54, 179–186. https://doi.org/10.1016/j.ympev.2009.08.023.

Rodríguez, R., 2015a. Nuevos registros de distribución de *Gambusia rhizophorae* (Teleostei: Poeciliidae) en el archipiélago cubano. NOVITATES CARIBAEA 8, 144–148.

Rodríguez, R., 2015b. *Rivulus berovidesi*, a new killifish species (Teleostei: Rivulidae) from western Cuba. Zootaxa 3949. http://dx.doi.org/10.11646/zootaxa.3949.2.9.

Ronquist, F., Teleslenko, M., Van der Mark, P., Ayres, D.L., Darling, A., Hohna, S., Larget, B., Liu, L., Suchard, M.A., Huelsenbeck, J.P., 2012. MrBayes 3.2: Efficient Bayesian phylogenetic inference and model choice across a large model space. Syst. Biol. 61, 539–542. http://doi.org/10.1093/sysbio/sys029.

Rosen, D., Bailey, R., 1963. The poeciliidae fishes (Cyprinodontiformes), their structure, zoogeography, and systematics. Bull Am Mus Nat Hist 126, 1–176.

Rousset, F., 2008. genepop’007: a complete re-implementation of the genepop software for Windows and Linux. Mol Ecol Resour 8, 103–106. 10.1111/j.1471-8286.2007.01931.x.

Scribner, K.T., Avise, J.C., 1993. Cytonuclear genetic architecture in mosquitofish populations and the possible roles of introgressive hybridization. Molecular Ecology 2, 139–149. http://doi.org/10.1111/j.1365-294X.1993.tb00103.x.

Sly, N.D., Townsend, A.K., Rimmer, C.C., Townsend, J.M., Latta, S.C., Lovette, I.J., 2011. Ancient islands and modern invasions: disparate phylogeographic histories among Hispaniola’s endemic birds. Molecular Ecology 20, 5012–5024. http://doi.org/10.1111/j.1365-294X.2011.05073.x.

Spencer, C.C., Chlan, C.A., Neigel, J.E., Scribner, K.T., Wooten, M.C., Leberg, P.L., 1999. Polymorphic microsatellite markers in the western morquitofish, *Gambusia affinis*. Molecular Ecology 8, 157–168.

Stamatakis, A., 2006. RAxML-VI-HPC: maximum likelihood-based phylogenetic analyses with thousands of taxa and mixed models. Bioinformatics 22, 2688–2690. http://doi.org/10.1093/bioinformatics/btl446.

Stamatakis, A., Blagojevic, F., Nikolopoulos, D.S., Antonopoulos, C.D., 2007. Exploring new search algorithms and hardware for phylogenetics: RAxML meets the IBM Cell. J VLSI Sign Process Syst Sign Im 48, 271–286. http://doi.org/10.1007/s11265-007-0067-4.

Tajima, F., 1989. Statistical method for testing the neutral mutation hypothesis by DNA polymorphism. Genetics 123, 585–595.

Tatarenkov, A., Lima, S.M.Q., Earley, R.L., Berbel-Filho, W.M., Vermeulen, F.B.M., Taylor, D.S., Marson, K., Turner, B.J., Avise, J.C., 2017. Deep and concordant subdivisions in the self-fertilizing mangrove killifishes (*Kryptolebias*) revealed by nuclear and mtDNA markers. Biological Journal of the Linnean Society 122, 558–578. http://doi.org/10.1093/biolinnean/blx103.

Thompson, J.D., Higgins, D.G., Gibson, T.J., 1994. Clustal W: improving the sensitivity of progressive multiple sequence alignment through sequence weighting, position-specific gap penalties and weigth matrix choise. Nucleic Acid Research 22, 4673–4680.

Tobler, M., Kelley, J.L., Plath, M., Riesch, R., 2018. Extreme environments and the origins of biodiversity: adaptation and speciation in sulfide spring fishes. Molecular Ecology 27, 843–859. http://doi.org/10.1111/mec.14497.

van Gestel, J.P., Mann, P., Dolan, J.F., Grindlay, N.R., 1998. Structure and tectonics of the upper Cenozoic Puerto Rico–Virgin Islands carbonate platform as determined from seismic reflection studies. Journal of Geophysical Research 103, 30505–30530.

Van Oosterhout, C., Hutchinson, W., Wills, D., Shipley, P., 2004. MICRO-CHECKER: software for identifying and correcting genotyping errors in microsatellite data. Molecular Ecology Notes 4, 535–538. http://doi.org/10.1111/j.1471-8286.2004.00684.x.

Vieites, D.R., Wollenberg, K.C., Andreone, F., Köhler, J., Glaw, F., Vences, M., 2009. Vast underestimation of Madagascar’s biodiversity evidenced by an integrative amphibian inventory. PNAS 106, 8267–8272. http://doi.org/10.1073/pnas.0810821106.

Vines, T.H., Schluter, D., 2006. Strong assortative mating between allopatric sticklebacks as a by-product of adaptation to different environments. Proceedings of the Royal Society B: Biological Sciences 273, 911–916. http://doi.org/10.1098/rspb.2005.3387.

Watterson, G.A., 1975. On the number of segregating sites in genetical models without recombination. Theoretical Population Biology 7, 256–276. https://doi.org/10.1016/0040-5809(75)90020-9.

Weaver, P.F., Cruz, A., Johnson, S., Dupin, J., Weaver, K.F., 2016. Colonizing the Caribbean: biogeography and evolution of livebearing fishes of the genus Limia (Poeciliidae). Journal of Biogeography 43, 1808–1819. http://doi.org/10.1111/jbi.12798.

Weir, B.S., Cockerham, C.C., 1984. Estimating *F*-statistics for the analysis of population structure. Evolution 38, 1358–1370.

Wilk, R.J., Horth, L., 2016. A genetically distinct hybrid zone occurs for two globally invasive mosquito fish species with striking phenotypic resemblance. Ecology and Evolution 6, 8375–8388. http://doi.org/10.1002/ece3.2562.

Zane, L., Nelson, W.S., Jones, A.G., Avise, J.C., 1999. Microsate assesment of multiple âternity in natural populations of a live-bearing fish, *Gambusia holbrooki*. J Evol Biol 12, 61–99.

Zhang, J., Kapli, P., Pavlidis, P., Stamatakis, A., 2013. A general species delimitation method with applications to phylogenetic placements. Bioinformatics 29, 2869–2876. http://doi.org/10.1093/bioinformatics/btt499.

