## Supplemental Material for "Phylogeographic evidence that the distribution of cryptic euryhaline species in the *Gambusia punctata* species group in Cuba was shaped by the archipelago geological history"

**Supplementary material 1a.** Information about the individuals sampled for this study and accession numbers of the analyzed data sets.

| | | | | | | | | Genebank accession number | | | | | | |
|---|---|---|---|---|---|---|---|---|---|---|---|---|---|---|
| Species | Pop No. | ID[b] | Collection locality (River, province) | GPS | Salinity[c] | Micros (n)[d] | *cyt*b haplotypes[e] | cytb | COI | 12SRNA-tRNAval-16SRNA | 16SRNA-tRNALeu-ND1 | COII-tRNALys | COIII | Control Region |
| *Gambusia rhizophorae* | 1 | Lva | El Vajazal Lagoon, Pinar del Río | 22.4179, -84.3255 | B | 4 | G2,G3 | LR597092; LR597093 | | | | | | |
| | 2 | Cam1 | Camarones River, Pinar del Río | 22.4280, -84.2928 | F | 10 | G3 | LR597093 | | | | | | |
| | 3 | Ebla | Las Blancas estuary, Pinar del Río | 22.4649, -84.2807 | B | | G1,G2 | LR597091;LR597092 | LR596616 | LR597302 | LR597280 | LR597213 | LR597069 | LR597257 |
| | 4 | Mc | Mil Cumbres, Pinar del Río | 22.7807, -83.3861 | F | | G4,G5 | LR597094; LR597095 | LR596618 LR596619 | LR597304 LR597305 | LR597282 LR597283 | LR597215 LR597216 | LR597071 LR597072 | LR597259 LR597260 |
| | 5 | RSC | San Claudio River, Pinar del Río | 22.8426, -83.0106 | F | 5 | G6 | LR597096 | | | | | | |
| | 6 | Cab1 | Stream near Cabañas, Pinar del Río | 22.9754, -82.8921 | F | | G2,G6 | LR597092; LR597096 | | | | | | |
| | 7 | Cap | Capellanía River, Pinar del Río | 22.8584, -82.6793 | F | | G7,G8 | LR597097; LR597098 | | | | | | |
| | 8 | B1 | Baracoa River, La Havana | 23.0423, -82.5712 | B | 21 | G3,G11, G14,G15, B1, B2 | LR597093; LR597101; LR597104; LR597105; LR597106; LR597107 | | | | | | |
| | 9 | B2 | Baracoa River, La Havana | 23.0361, -82.5698 | F/B | 10 | G14, B1, B3 | LR597104; LR597106; LR597108 | | | | | | |
| | 10 | Gua | Guanabo River, La Havana | 23.1656, -82.1221 | B | 5 | G9 | LR597099 | | | | | | |
| | 11 | Bjar | Boca de Jaruco, La Havana | 23.1672, -82.0014 | F | 5 | G10,G11 | LR597100; LR597101 | | | | | | |
| | 12 | Ji | Jibacoa River (km 64), La Havana | 23.1315, -81.7964 | F | 6 | G12,G13 | LR597102; LR597103 | LR596617 | LR597303 | LR597281 | LR597214 | LR597070 | LR597258 |
| *Gambusia punctata* | 13 | Pap | Papaya River, Pinar del Río | 22.2493, -83.8465 | F | 3 | B14,B20, B21 | LR597119; LR597125; LR597126 | LR596620 | LR597306 | LR597284 | LR597217 | LR597073 | LR597261 |
| | 14 | Con | El Convento River, Pinar del Río | 22.3519, -83.3953 | F | | B14 | LR597119 | LR596621 | LR597307 | LR597285 | LR597218 | LR597074 | LR597262 |
| | 15 | Plc | Los Palacios River, Pinar del Río | 22.6126, -83.2816 | F | | B18,B19 | LR597123; LR597124 | | | | | | |
| | 16 | SC | San Cristóbal, Pinar del Río | 22.6995, -83.0203 | F | | B24,B25, B26 | LR597129; LR597130; LR597131 | | | | | | |
| | 17 | AMas | Masón Stream, Pinar del Río | 22.8468, -82.9741 | F | | B22,B23 | LR597127; LR597128 | | | | | | |
| | 18 | SJ | San Juan River, Pinar del Río | 22.8559, -82.9423 | F | 10 | B16,B17 | LR597121; LR597122 | | | | | | |
| | 19 | Gu | Guanimar La Havana | 22.6945, -82.6516 | B | 9 | B1,B8 | LR597106; LR597113 | | | | | | |
| | 20 | HabP | Havana-Pinar del Río Road | 22.8625, -82.8589 | F | | B1,B5 | LR597106 | | | | | | |
| | 21 | APNB | Stream Niña Bonita dam, Pinar del Río | 23.0112, -82.5079 | F | | B1 | LR597106 | | | | | | |
| | 22 | Yag | Ariguanabo River La Havana | 22.8979, -82.4951 | F | | B1,B4 | LR597106; LR597109 | | | | | | |
| | 23 | Gr | La Grifa Key, La Broa Cove, La Havana | 22.2304, -82.8170 | S | | B11 | LR597116 | LR596622 | LR597308 | LR597286 | LR597219 | LR597075 | LR597263 |
| | 24 | Ita | Itabo River, I. Juventud | 21.6889, -82.9787 | F | 5 | - | | | | | | | |

| | | | | | | | | | | | | | | |
|---|---|---|---|---|---|---|---|---|---|---|---|---|---|---|
| | 25 | Scan | La Cañada, I. Juventud | 21.7220, -82.9075 | F | | B15 | LR597120 | | | | | | |
| | 26 | Sigua | La Siguanea, I. Juventud | 21.5970, -82.9185 | B | | B12,B13, B14 | LR597117; LR597118; LR597119 | LR596623 LR596624 | LR597309 LR597310 | LR597287 LR597288 | LR597220 LR597221 | LR597076 LR597077 | LR597264 LR597265 |
| | 28 | Charp | Palpite Marsh, Matanzas | 22.3337, -81.1816 | F | | B1,B5,B6 | LR597106; LR597110; LR597111 | | | | | | |
| | 29 | Pal | Palpite, Matanzas | 22.3323, -81.1812 | F | 10 | B10 | LR597115 | | | | | | |
| | 30 | Lfac | Facundo Lagoon, Matanzas | 22.2794, -81.1632 | F | 4 | B9 | LR597114 | | | | | | |
| | 31 | CPat | Los Patos Chanel, Matanzas | 22.2528, -81.1481 | F | 3 | B1 | LR597106 | | | | | | |
| | 32 | Csag | Cárdenas-Sagua Grande,Matanzas | 22.9093, -81.0546 | F | 4 | B1 | LR597106 | | | | | | |
| | 33 | CaSab | Cárdenas-Sagua Grande,Villa Clara | 22.9776, -80.6645 | F | | B1,B7 | LR597106; LR597112 | LR596625 | LR597311 | LR597289 | LR597222 | LR597078 | LR597266 |
| *Gambusia* sp. D[a] | 27 | Cani | Canimar River, Matanzas | 23.0368, -81.4921 | F | | O2,O3 | LR597133; LR597134 | LR596626 | LR597312 | LR597290 | LR597223 | LR597079 | LR597267 |
| | 34 | Ranch | River near Ranchuelos, Villa Clara | 22.4058, -80.1620 | F | 5 | O1 | LR597132 | LR596627 | LR597313 | LR597291 | LR597224 | LR597080 | LR597268 |
| | 35 | Ari | Ariamo River, Cienfuegos | 22.0527, -80.2919 | F | 4 | O8 | LR597139 | | | | | | |
| | 36 | Gav | Gavilanes River, Cienfuegos | 21.9888, -803206 | F | | O11,O12 | LR597142 ; LR597143 | LR596628 | LR597314 | LR597292 | LR597225 | LR597081 | LR597269 |
| | 37 | Rjua | San Juan River, Cienfuegos | 21.9189, -80.2660 | F | | O13,O14 | LR597144 ; LR597145 | LR596629 | LR597315 | LR597293 | LR597226 | LR597082 | LR597270 |
| | 38 | Rhon | Hondo River, Cienfuegos | 21.8327, -80.1504 | F | | O6,O7 | LR597137 ; LR597138 | LR596630 | LR597316 | LR597294 | LR597227 | LR597083 | LR597271 |
| | 39 | Caba | Cabagán River,Cienfuegos/V.Clara | 21.8252, -801135 | F | | O4,O5 | LR597135 ; LR597136 | | | | | | |
| | 40 | Bta | Batata Stream, Cienfuegos | 21.8973, -80.0441 | F | 9 | O15 | LR597146 | | | | | | |
| | 41 | Cabu | El Caburní, Sancti Spiritus | 21.9279, -80.0360 | F | 4 | O10 | LR597141 | | | | | | |
| | 42 | 042 | Yaguanabo River, Sancti Spiritus | 21.8117, -80.0008 | F | 9 | O5,O9 | LR597136; LR597140 | | | | | | |
| | 43 | Agab | Agabama River, Villa Clara | 22.2643, -79.8600 | F | 4 | O5 | LR597136 | LR596631 | LR597317 | LR597295 | LR597228 | LR597084 | LR597272 |
| *Gambusia* sp. D + *Gambusia* sp. hybrids | 44 | Cal | Calabazas River, Sancti Spiritus | 22.1697, -79.5717 | F | | R5,R6 | LR597151 ; LR597152 | LR596632 | LR597318 | LR597296 | LR597229 | LR597085 | LR597273 |
| | 45 | Hig | Higuanojo River, Sancti Spiritus | 21.7975, 79.6794 | F | 5 | R1,R2 | LR597147 ; LR597148 | | | | | | |
| | 46 | Tay | Tayabacoa River, Sancti Spiritus | 21.8046, -79.5857 | F | | R1,R13 | LR597147 ; LR597159 | | | | | | |
| | 47 | Cay | Cayajana River, Sancti Spiritus | 21.8600, -79.5126 | F | | R16,R17 | LR597162; LR597163 | | | | | | |
| | 48 | ZM | Zaza de Medio, Sancti Spiritus | 21.9961, -79.3669 | F | 5 | R1 | LR597147 | | | | | | |
| | 49 | Tag | La Alforja, Sancti Spiritus | 21.9913, -79.2530 | F | | R1,R21 | LR597147 ; LR597167 | | | | | | |
| | 50 | Jat | Jatibonico River, Sancti Spiritus | 21.9487, -79.1759 | F | | R14,R15 | LR597160; LR597161 | LR596633 | LR597319 | LR597297 | LR597230 | LR597086 | LR597274 |
| | 51 | Rmaj | Majagua River, Ciego de Avila | 21.9324, -78.9994 | F | 5 | R1,R13 | LR597147 ; LR597159 | LR596634 | LR597320 | LR597298 | LR597231 | LR597087 | LR597275 |
| | 52 | Cor | Corrales River, Ciego de Avila | 21.8979, -78.9324 | F | | R1,R12 | LR597147 ; LR597158 | | | | | | |
| | 53 | Cjen | La Jenifer sinkhole, Ciego de Avila | 22.5373, -78.4072 | B | | R3 | LR597149 | | | | | | |
| | 54 | Sole | Soledad River, Ciego de Avila | 21.6755, -78.3868 | F | | R1 | LR597147 | | | | | | |
| | 55 | Jrc | Cuervo Key, Ciego de Avila | 21.0623, -78.9632 | S | | R19,R20 | LR597165; LR597166 | | | | | | |
| | 56 | Gra | Algodón Grande Key, Ciego de Avila | 21.1090, -78.7357 | S | 6 | R13,R18 | LR597159; LR597164 | LR596635 | LR597321 | LR597299 | LR597232 | LR597088 | LR597276 |
| | 57 | SAna | Santa Ana Stream, Camagüey | 21.3054, -78.3724 | F | | R1 | LR597147 | | | | | | |
| | 58 | SMar | Santa María River, Camagüey | 21.4004, -78.0593 | F | 10 | R1 | LR597147 | LR596636 | LR597322 | LR597300 | LR597233 | LR597089 | LR597277 |

| | | | | | | | | | | | | | | |
|---|---|---|---|---|---|---|---|---|---|---|---|---|---|---|
| | 59 | Braz | El Brazo River, Las Tunas | 21.1392, -77.6133 | F | | R7,R11 | LR597153 ; LR597157 | | | | | | |
| | 60 | Joba | Jobabo River, Las Tunas | 20.9155, -77.3039 | F | 10 | R4 | LR597150 | | | | | | |
| *Gambusia* sp. | 61 | Gib | Cerro Colorado Stream, Holguín | 21.0965, -76.1517 | F | 5 | R4 | LR597150 | | | | | | |
| | 62 | Ryar | Yara River, Gramma | 20.3433, -77.0615 | F | 11 | R7 | LR597153 | LR596637 | LR597323 | LR597301 | LR597234 | LR597090 | LR597278 |
| | 63 | Bay | Bayamo River, Gramma | 20.3650, -76.6496 | F | 10 | R4,R22 | LR597150; LR597168 | | | | | | |
| | 64 | Yar | Yara River, Gramma | 20.1198, -76.9236 | F | | R7,R8 | LR597153 ; LR597154 | | | | | | |
| | 65 | Rcoj | Cojímar River, Santiago de Cuba | 19.9710, -76.1012 | F | 10 | R9,R10 | LR597155; LR597156 | | | | | | |
| | 66 | Cua | Las Cuabas, Santiago de Cuba | 20.0515, -75.8103 | F | 2 | R1 | LR597147 | | | | | | |
| *G. rhizophorae*[f] | - | - | Key West, Florida | - | - | - | - | U18223 | - | - | - | - | - | - |
| *G. rhizophorae*[g] | - | - | Matheson Hammock, Florida | - | - | - | - | KM658368 | - | - | - | - | - | - |
| *Gambusia puncticulata* C[h] (out group) | - | - | Cañete River, Guantánamo | 20.6227, -74.8529 | F | | | LR597427 | LR597428 | LR597429 | LR597430 | LR597431 | LR597432 | LR597433 |
| *G. hispaniolae*[f] | - | - | Dominican Republic | - | - | - | - | U18209 | - | - | - | - | - | - |

[a] : A redefinition of *Gambusia* sp. in Lara et al. (2010) (see text).
[b]: Acronym used for each locality.
[c]: F, freshwater (<0.5ppt); B, brackish water (0.5 – 30 ppt); S, salt water (30 – 50ppt).
[d]: Number of exemplars analyzed for microsatellite loci at each locality.
[e]: Haplotype label prefixes (B, bleu clade; G, green clade; O, orange clade; R, red clade) equate clades colors in Fig. 2 and Supplementary material 2.
[f]: Lydeard et al. (1995)
[g]: Heinen-Kay et al. (2014)
[h]: *Gambusia puncticulata* C sensu Lara et al. (2010).

**Supplementary material 1b**: Amplification and sequencing primers used in the study of *G. punctata* species group.

| Primer name | Gene segment | Primer sequence 5’ -> 3’ | Sequence Length (bp) | Source |
|---|---|---|---|---|
| GluGamb<br>CB3 | *cyt*b | ACT CAA CTA TAA GAA CYC TAA TGG C<br>TGC GAA GAG GAA GTA CCA TTC | 752 | Meyer et al. (1990)*<br>Palumbi, 1996 |
| 12S-L1090<br>16S-H1782 | 12SRNA – tRNAVal - 16SRNA | AAA CTG GGA TTA GAT ACC CCA CTA<br>TTT CAT CTT TCC CTT GCG GTA C | 571 | Hrbek and Larson, 1999 |
| 16S-L3079<br>tRNAIle-H4280 | 16SRNA-tRNALeu-ND1 | ACG TGA TCT GAG TTC AGA CCG<br>ACT GTA TCA AAG TGG YCC TT | 1166 | Hrbek and Meyer, 2003 |
| FishCOIf<br>FishCOIr | COI | AAY CAY AAA GAY GGY ACC CT<br>CNG GRT GN C CRA AGA AYC | 648 | Lara et al. 2010 |
| GambCOIIf<br>GambLysAR | COII-tRNALys | CAA CTA GGY TTT CAA GAT GC<br>CMA TTT TTA GCT TAA AAG GC | 669 | This study |
| CO3Gamb2F<br>GambCO3R | COIII | CAA GCA CAT GCA TAT CAY ATA<br>ACR TCR ACG AAA TGT CAR TAT CA | 649 | This study |
| CR-F<br>CR-B | Control region | ATG CCA GGA ATA GTT CAC CGT GT<br>TGT ATG TAT TAT CCC CAT TAA TCT | 308 | This study |

* Modified

Supplementary material 2a.

Maximum likelihood phylogenetic tree (LogL = - 2793.97) based on 752 bp cytb sequences. The model Tamura-Nei 1993 with gamma correction (α = 0.14) and invariants (I = 0.58) were used. ML bootstrap and Bayesian posterior probabilities, in that order, are represented for each node. Zero values indicates the node is absent on the ML or the Bayesian phylogenetic trees.

Rate matrice: 1.0000, 12.6303, 1.0000, 1.0000, 6.2786, 1.0000
Base frequencies: A=0.2306, T=0.3200, G=0.1565, C=0.2929

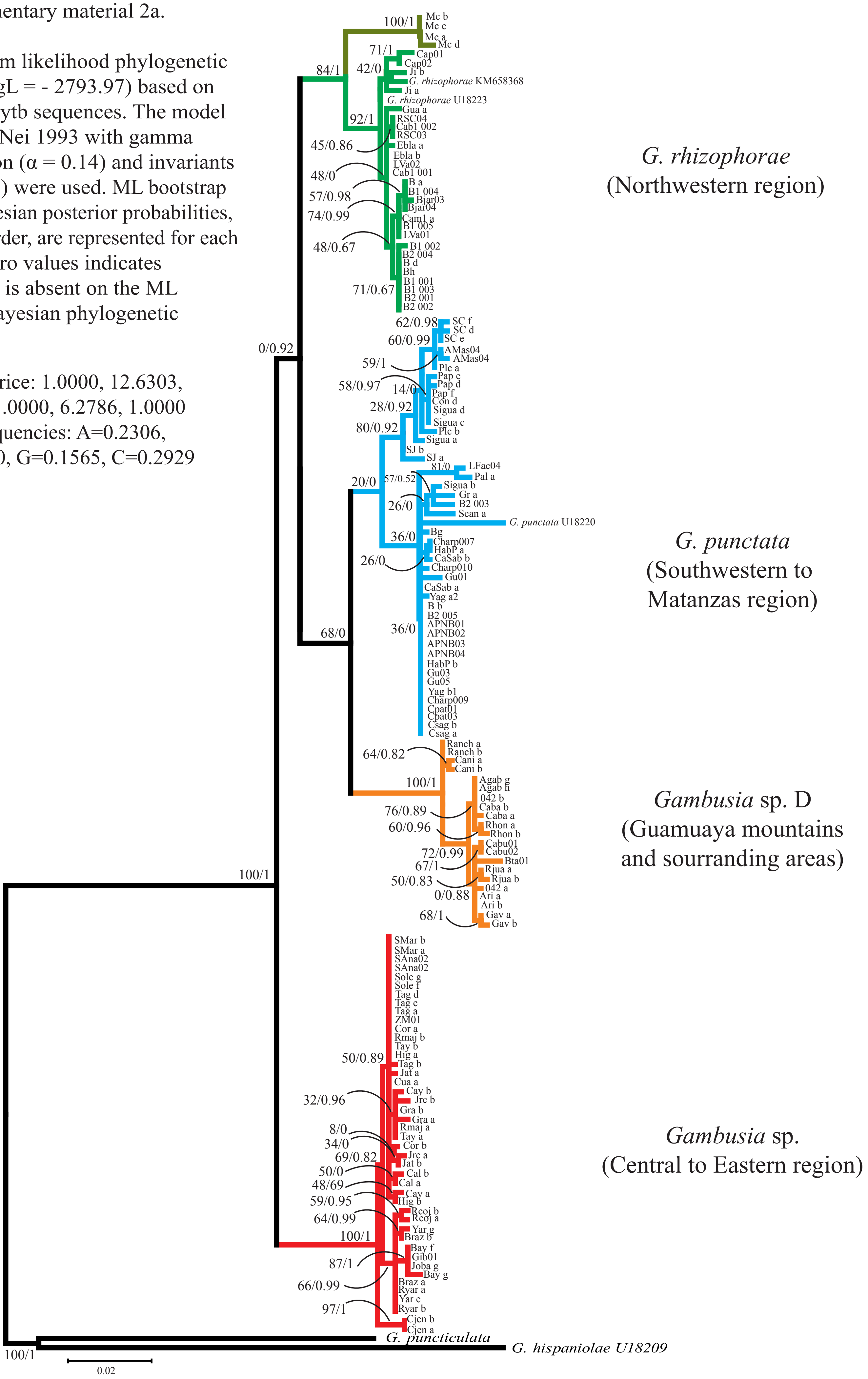

Supplementary material 2b. Maximum likelihood phylogenetic tree (LogL = - 12366.47) based on 4,763 bp of cytb, 12SRNA - tRNAVal - 16SRNA, 16SRNA - tRNALeu - ND1, COI, COII -tRNALys, COIII and control region partial sequences. TheTamura-Nei (1993 ) model with gamma correction (α = 0.15) was used. ML bootstrap and Bayesian posterior probabilities, in that order, are represented for each node.

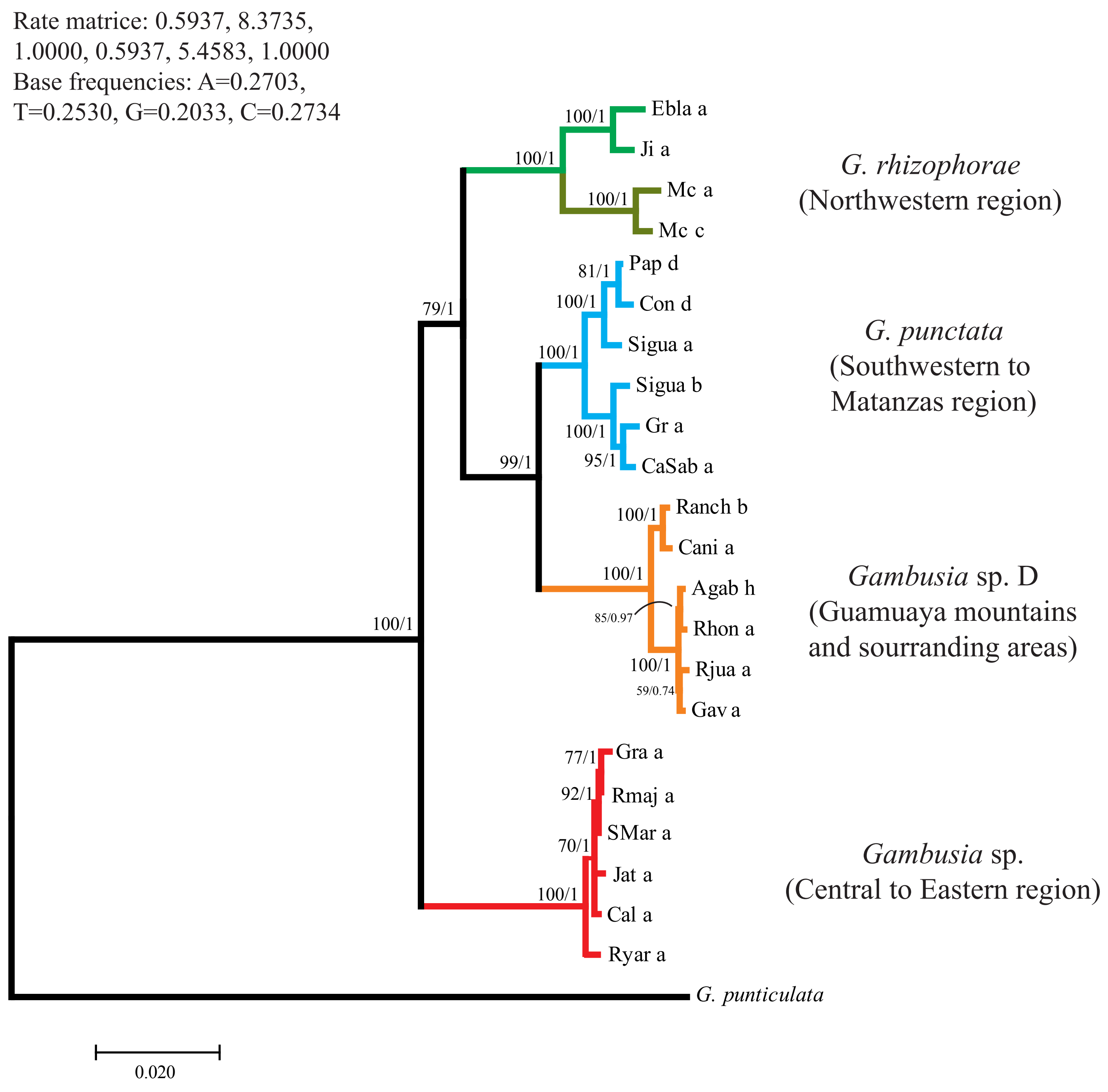

Supplementary material 2c.

Bayesian phylogenetic tree based on 752 bp cytb sequences. The model Tamura-Nei 1993 with gamma correction (α = 0.14) and invariants (I = 0.58) were used. Bayesian posterior probabilities, higher than 0.95 are represented for each node.

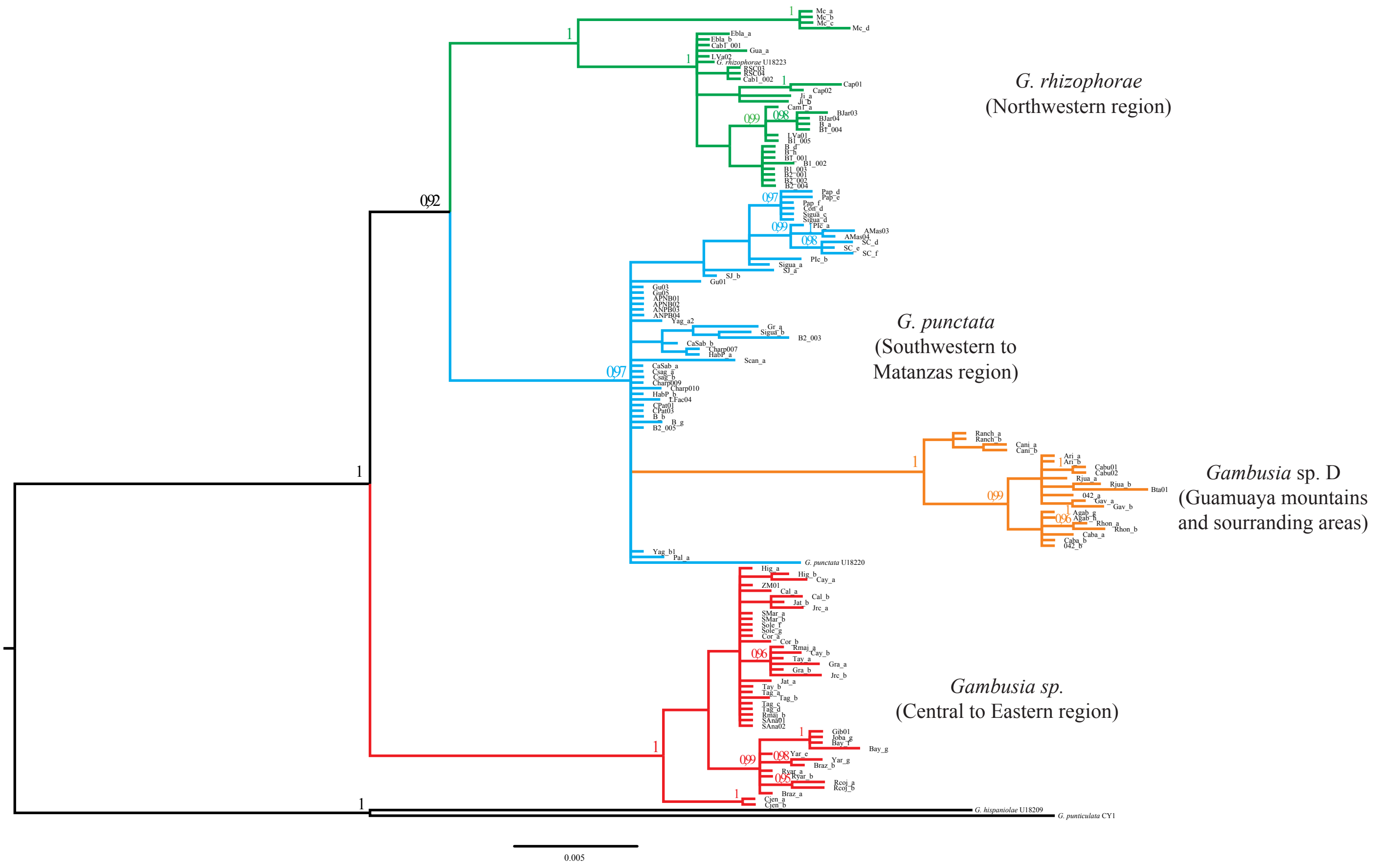

**Supplementary material 3**

Results of the Poisson tree processes (PTP) model analysis based on *cyt*b and concatenated data set ML trees. We used a RAxML (Stamatakis 2006) tree as the input obtained using the GTRCAT substitution model. Bayesian support values are shown for the main partitions recovered.

| Partition | Concatenate | *cyt*b |
|---|---|---|
| *G. rhizophorae* sensu stricto | 0.817 | 0.633 |
| *G. rhizophorae* (Mil Cumbres) | 0.784 | 0.615 |
| *G. punctata* | 0.597 | - |
| *Gambusia* sp. D | 0.859 | 0.567 |
| *Gambusia* sp. | 0.976 | 0.300 |

RAxML (Stamatakis 2006) trees used as the input. A) cytb sequences; B) Concatenated data set.

A)

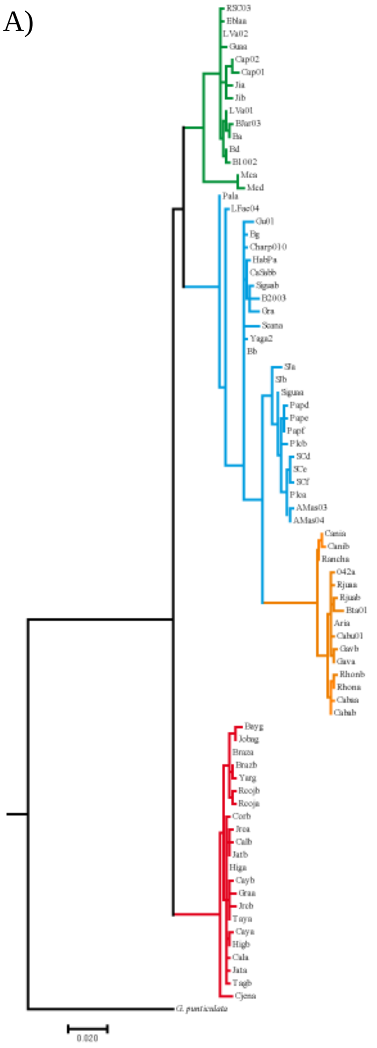

B)

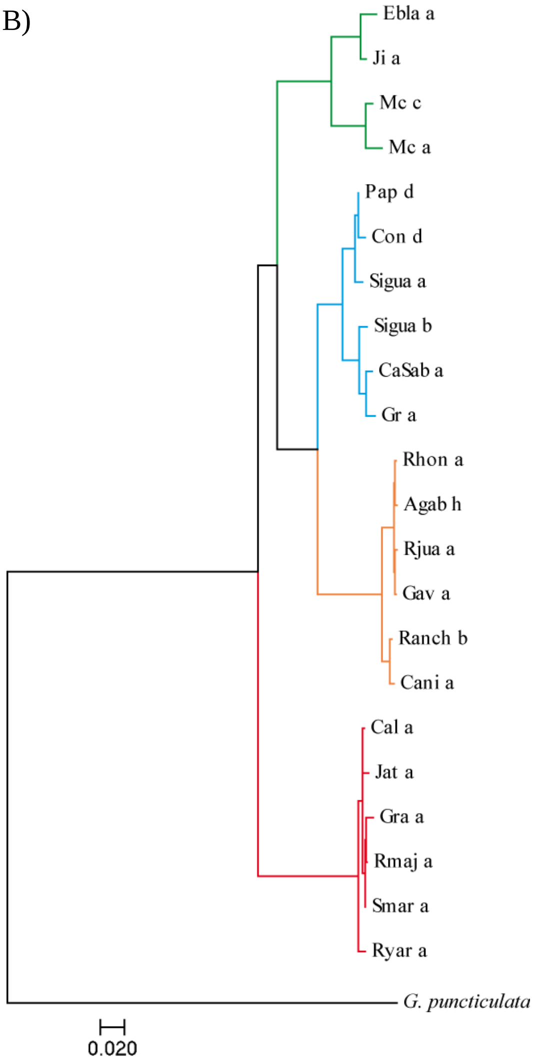

### Supplementary Material 4

Hardy – Weinberg test with $F_{IS}$ estimates and probabilities (among brackets) per locus and population. Statistically significant departures from equilibrium frequencies (after Bonferroni correction p<0.001) are in bold. Estimates were obtained by the probability test option using Markov chain method with 10000 dememorizations, 20 batches and 5000 iterations per batch.

| | | ***Locus*** | | | | | | | | |
|---|---|---|---|---|---|---|---|---|---|---|
| **Species** | **Locality** | ***Gaaf* 10** | ***Gaaf* 13** | ***Gaaf* 15** | ***Gaaf* 16** | ***Gaaf* 22** | ***Gafμ* 5** | **GG2B** | ***Mf* 13** | ***Mf* 6** |
| ***G. rhizophorae*** | Lva | 0.182 (0.409) | 0.250 (0. 419) | 1.000 (0.140) | 0.700 (0.029) | 0.000 (0.308) | -0.200 (1.000) | 0 | -0.500 (1.000) | 0 |
| | Cam | 0.361 (0.037) | 0.379 (0.105) | 0.022 (0.529) | **0.316 (0.000)** | 0.105 (0.638) | 0.106 (0.431) | 0 | -0.145 (0.108) | -0.125 (1.000) |
| | RSC | 0 | 0 | -0.600(0.428) | 1.000 (0.111) | -0.538 (0.430) | 0.273 (1.000) | 0.600 (0.336) | 0 | 0 |
| | B1+B2 | 0.324 (0.037) | -0.085 (1.000) | -0.041 (0.520) | 0.303(0.147) | 0.005 (0.781) | 0.034 (0.712) | 0.161 (0.624) | 0.078 (0.274) | 0.048 (0.469) |
| | Gua | 0.733 (0.046) | 1.000 (0.049) | 0 | 1.000 (0.009) | -0.143 (1.000) | 0 | -0.143 (1.000) | -0.429 (0.747) | -0.230 (1.000) |
| | Bjar | 0.030 (0.846) | -0.143 (1.000) | -0.143 (1.000) | 0.077 (1.000) | 0.733 (0.047) | 0.483 (0.547) | 0 | -0.250 (0.902) | 0 |
| | Ji | 0.000 (0.720) | 0 | 0 | 0.615 (0.271) | 0 | 0 | 0 | 0.706 (0.155) | 0 |
| ***G. punctata*** | Pap | -0.091 (1.000) | 0.000 (1.000) | -0.333 (1.000) | 0.111 (1.000) | 0.273 (0.479) | -0.333 (1.000) | 0 | -0.200 (1.000) | 0 |
| | SJ | 0 | 0 | 0 | -0.046 (1.000) | 0 | 0 | 0 | -1.000 (0.007) | 0 |
| | Gu | 0.103 (0.958) | 0.238 (0.089) | -0.217 (1.000) | -0.131 (0.917) | 0.213 (0.205) | 0.123 (1.000) | 0 | 0.121 (0.642) | -0.231 (1.000) |
| | Ita | -0.081 (1.000) | 0.111 (0.610) | -0.143 (1.000) | 0 | 0.059 (0.839) | 0 | 0 | 0.172 (1.000) | 0.636 (0.113) |
| | Cpat+Pal+Lfag | 0.006 (0.770) | 0.118 (0.082) | -0.096 (1.000) | -0.103 (1.000) | 0.107 (0.883) | 0.285 (0.289) | 0 | 0.053 (0.304) | 0.074 (0.308) |
| | Csag | 0.250 (0.445) | 0.000 (1.000) | 0 | 0 | -0.412 (1.000) | 0.286 (1.000) | 0 | -0.412 (1.000) | 0.571 (0.430) |
| ***Gambusia* sp. D** | Ranch | -0.333 (1.000) | 0 | 0 | 0 | 0.250 (0.697) | -0.391 (1.000) | 0 | 1.000 (0.110) | 0 |
| | Ari | 0 | -0.125 (1.000) | 0 | 0 | 1.000 (0.143) | 0 | -1.000 (0.313) | 0 | 0 |
| | Bta | 0 | -0.111 (1.000) | 0 | 0 | 1.000 (0.076) | 0.636 (0.178) | 0 | 0.407 (0.341) | 0 |
| | 042 | -0.103 (0.328) | -0.032 (1.000) | 0 | 0 | 0.199 (0.156) | 0 | 0 | 0.121 (0.633) | 0 |
| | Cabu | 0 | -0.500 (1.000) | 0 | 0 | -0.200 (1.000) | -0.200 (1.000) | 0 | 0 | -1.000 (0.312) |
| | Agab | 0.217 (0.311) | 0 | 0 | 0 | 0.368 (0.304) | 0 | 0 | 0.294 (1.000) | 0 |
| **Hybrid zone** | Hig | 0.143 (0.613) | 0.000 (1.000) | 0 | 0 | -0.125 (1.000) | 0 | 0 | 0 | 0 |
| | ZM | 0.077 (1.000) | -0.600 (0.430) | 0 | 0 | -0.143 (1.000) | 0 | -0.333 (1.000) | 0 | 0 |
| | Rmaj | 0.111 (0.597) | 0.000 (0.618) | 0 | 0.273 (1.000) | 0 | 0 | 0 | -0.091 (1.000) | 0 |
| | Gra | 0.130 (0.399) | -0.136 (0.351) | 0 | 0 | 0.388 (0.105) | -0.111 (1.000) | 0 | 0.259 (0.273) | 0 |
| | Smar | 0.379 (0.153) | 0.343 (0.192) | 0 | -0.113 (1.000) | -0.036 (0.814) | 0.653 (0.052) | 1.000 (0.053) | -0.136 (1.000) | 0 |
| | Joba | 0.444 (0.004) | 0.337 (0.029) | 0 | 0.073 (0.393) | 0.238 (0.101) | 0.395 (0.105) | 0.100 (0.100) | 0.225 (1.000) | 0 |

| | | | | | | | | | | |
|---|---|---|---|---|---|---|---|---|---|---|
| ***Gambusia* sp.** | Gib | 0.143 (0.112) | 0.000 (0.648) | 0 | 0 | -0.067 (1.000) | -0.333 (1.000) | -0.143 (1.000) | 0.407 (0.244) | 0 |
| | Ryar | -0.149 (0.859) | -0.023 (0.424) | 0 | -0.071 (1.000) | -0.233 (0.910) | 0.642 (0.143) | -0.333 (0.505) | 0.411 (0.229) | 0 |
| | Bay | 0.207 (0.148) | 0.261 (0.008) | 0 | 0.077 (0.088) | -0.025 (1.000) | 0.454 (0.107) | 1.000 (0.052) | 0.189 (0.772) | -0.125 (1.000) |
| | Rcoj | -0.231 (1.000) | 0.400 (0.168) | 0 | 1.000 (0.053) | 0.755 (0.001) | 0 | 0.640 (0.157) | 0.724 (0.013) | 0 |
| | Cua | 0.500 (0.334) | 0.500 (0.344) | 0 | 0 | 1.000 (0.332) | -1.000 (1.000) | 0 | 0 | 0 |

0: All individuals where homozygous for a single allele.

**Supplementary material 5a**

The result of the Evanno's analysis (Evanno et al., 2005) indicating *K* = 4 as truth number of clusters. The analysis was performed with STRUCTURE HARVESTER v.0.6 (Earl and vonHoldt, 2012) using 20 simulations set of posterior individual membership probabilities for *K* = 2 to *K* = 7.

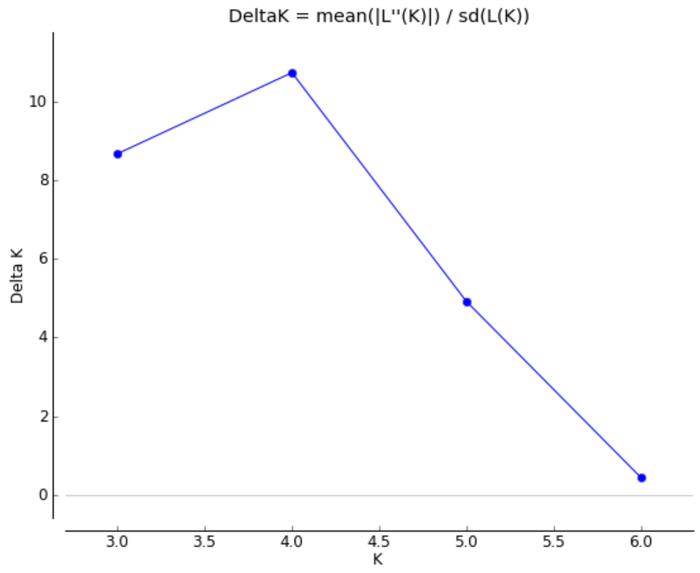

The plot of likelihoods of *K* (Ln Pr(X|K)).

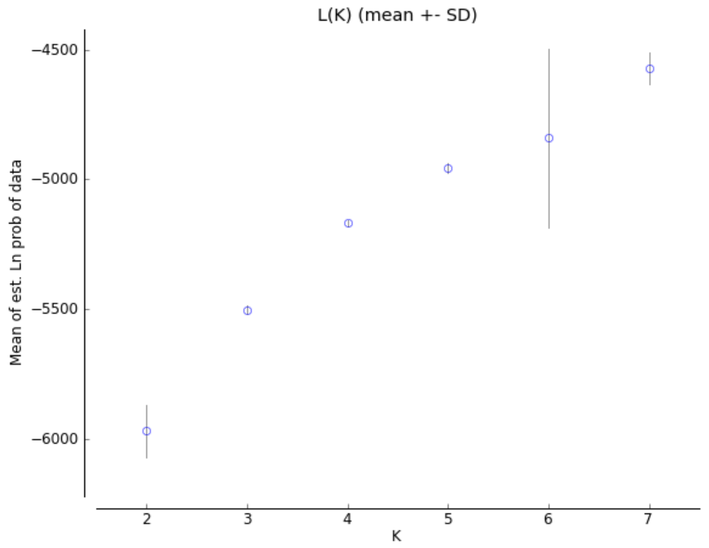

**Supplementary material 5b**

The results of the Evanno’s analysis (Evanno et al., 2005) conducted for the different lineages. The analysis was performed with STRUCTURE HARVESTER v.0.6 (Earl and vonHoldt, 2012) using 20 simulations set of posterior individual membership probabilities. Different *K* ranges were set according to the group.

Plots of likelihoods of *K* (Ln Pr(X|K))

*K* = 1 to *K* = 8

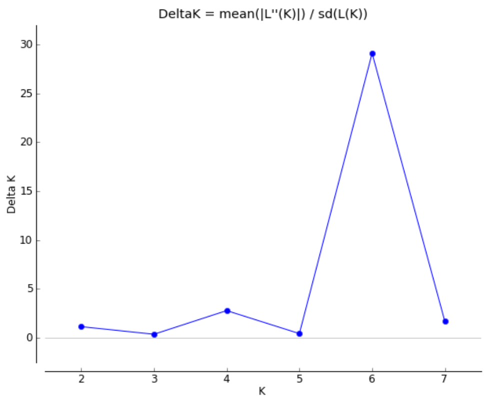

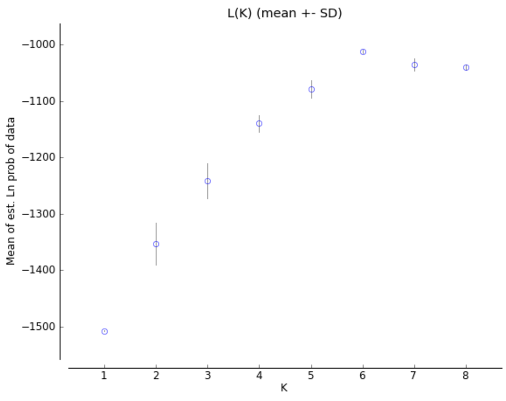

*Gambusia rhizophorae*

*K* = 1 to *K* = 5

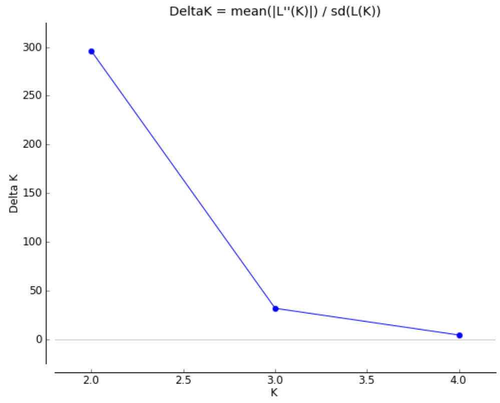

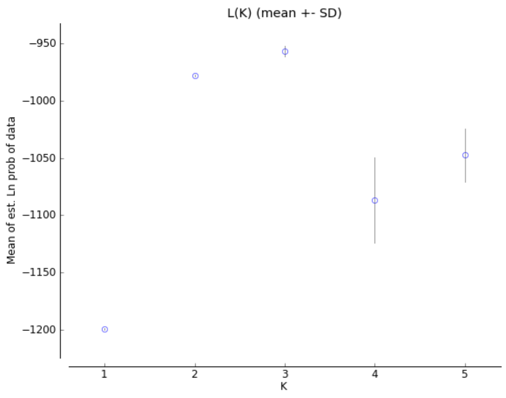

*Gambusia punctata*

*K* = 1 to *K* = 5

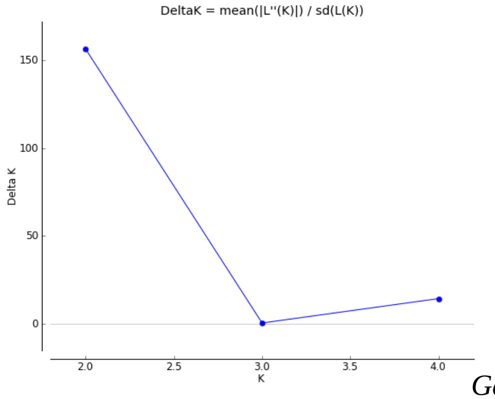

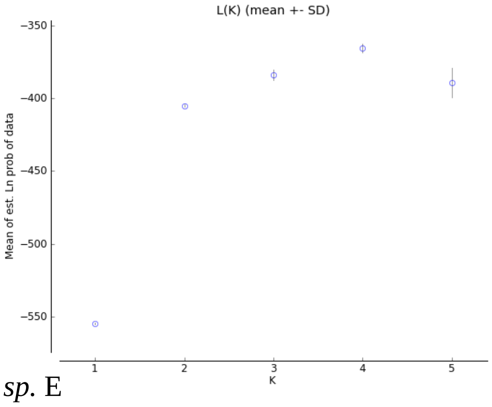

*Gambusia sp.* E

Plots of likelihoods of *K* (Ln Pr(X|K))

*K* = 1 to *K* = 5

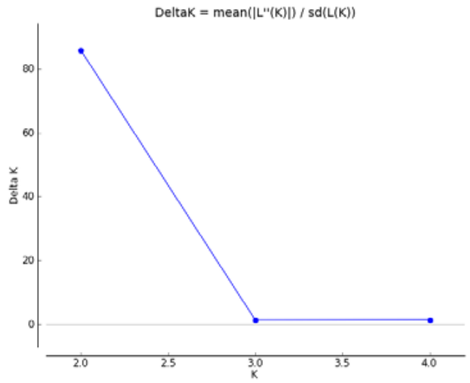

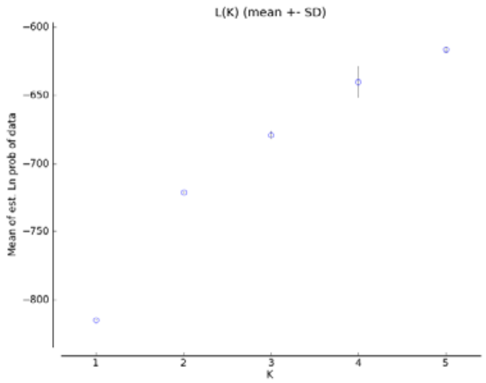

*Gambusia sp.*

**Supplementary material 6**. *Cyt*b gene diagnostic combinations and private DNA positions for the different species of the *Gambusia punctata* species group in Cuba. Localities from the hybrid zone were excluded. Lowercase indicates nucleotide frequencies below 5% in the population. Mc: Mil Cumbres population. [a]: *G. rhizophorae* populations excluding Mil Cumbres.

| | Species (Group) | | | | |
|---|---|---|---|---|---|
| Nucleotide position | *G. rhizophorae* | | *G. punctata* | *Gambusia* sp. D | *Gambusia* sp. |
| | All[a] | Mc | | | |
| 13 | C | T | C | C | C |
| 25 | A | A | A/G | C | A/G |
| 64 | C | C | T/c | T | T |
| 76 | T | T | T | C | T |
| 79 | A | A | A | G | A |
| 82 | G | A | G | G | G |
| 95 | G | G | G | G | A |
| 104 | C | C | C | C | T |
| 167 | A | A | G | G | G |
| 175 | T | T | T | C | T |
| 211 | T | T | C | C | C |
| 232 | T | T | T | C/t | T |
| 250 | C | T | T | T | T |
| 253 | C | C | C | C/T | C |
| 325 | T | T | C | T | T |
| 328 | T | C | C/T | T | T |
| 332 | C | C | C | C | T |
| 334 | G | A | G | G | A |
| 337 | G/A | A | A | A | A |
| 343 | G | G | A | A | A |
| 376 | A | A | A | G | A |
| 391 | A | A | A | A | G |
| 412 | T | C | T/C | T | T |
| 427 | G | G | G | A | G |
| 487 | C /t | C | C | C | T |
| 514 | C | T | C | C | C |
| 547 | C | C | C | T | C |
| 559 | T | T | T | A | T |
| 562 | T | T | T | T | C |
| 568 | T | T | T | T | C |
| 574 | G | G | G | A | G |
| 589 | C | C | T/c | T | T |
| 598 | A | A | A | G | A |
| 607 | C | C | C | T | C |
| 608 | G | G | G | G | A |
| 637 | C | T | C | C | C |
| 673 | C | C | C | C | A |
| 680 | G | G | A | G | G |
| 685 | A | A | G/A | G | G |
| 691 | C | T | T/c | T | T |
| 706 | C | C | T/C | T | C |
| 721 | G | A | G/A | G | G |
| 727 | A | C | A | A | A |
| 751 | C | C | C | T | C |
